# CRISPR activation of endogenous *PKD1* increases polycystin-1 levels and suppresses cellular features of ADPKD

**DOI:** 10.64898/2026.05.23.727418

**Authors:** Anubhav Chakraborty, Meekha M. Varghese, Darren P. Wallace, Christopher J. Ward, Alan S.L. Yu

## Abstract

Most cases of autosomal dominant polycystic kidney disease (ADPKD) are caused by mutations in *PKD1,* which reduce polycystin-1 (PC1) levels below a critical functional threshold. Normalizing PC1 dosage mitigates disease progression; therefore, we sought to develop a CRISPR activation (CRISPRa) strategy to transcriptionally upregulate endogenous *PKD1*. We systematically screened multiple single-guide RNAs using an EGFP-reporter platform and identified potent candidates targeting the proximal *PKD1* promoter in mouse and human cell models. Our results demonstrate that CRISPRa effectively increased endogenous *Pkd1* mRNA in the mouse collecting duct-derived *Pkd1^RC/-^*cell model and in the primary renal epithelial cells from PKD mice. In *Pkd1^RC/-^* cells, CRISPRa of *Pkd1* increased PC1 protein levels and significantly reduced cell proliferation and *in vitro* cyst formation in 3D cultures. Mechanistically, *Pkd1* activation improved mitochondrial membrane potential, reduced dependency on aerobic glycolysis, and corrected signaling pathways involved in cystogenesis, specifically reducing intracellular cAMP, cMyc, pCreb, and pErk levels, while increasing pYap1 levels. We confirmed the translational potential of this platform by successfully activating *PKD1* in primary renal epithelial cells from human kidneys. We observed a heterogeneous response across both normal and ADPKD patient-derived donor lines, with significant upregulation achieved in two of the tested cell preparations. These findings provide a compelling proof-of-concept that CRISPRa-mediated gene augmentation can increase PC1 levels, establishing a foundation for promising gene therapies aimed at successfully suppressing the pathogenic features of ADPKD.

**New and noteworthy:** This study provides proof-of-concept for a CRISPR activation (CRISPRa)-based approach to ADPKD. CRISPRa targeting the *PKD1* promoter, increased polycystin-1 levels in mouse and human cells. Upregulation of a hypomorphic *Pkd1* allele increased functional polycystin-1, corrected dysregulated signaling pathways, suppressed cell proliferation and *in vitro* cyst formation. These results establish CRISPRa as a promising therapeutic approach to restore polycystin levels above a critical threshold and suppress cystogenesis in ADPKD.

## Introduction

Autosomal dominant polycystic kidney disease (ADPKD) is the most common inherited monogenic disorder with an estimated prevalence of 1 in 1,000 individuals and the leading genetic cause of end-stage kidney disease worldwide (1). It is characterized by progressive development of fluid-filled renal cysts that expand over time, replacing the normal parenchyma and compromising normal renal architecture. This relentless cyst growth results in massive bilateral kidney enlargement, chronic pain, hypertension, and ultimately kidney failure (2). Most ADPKD cases are caused by mutations in *PKD1* (approximately 80%) and *PKD2* (about 15%) (2). *PKD1* and *PKD2* encode the integral membrane proteins, polycystin-1 (PC1) and polycystin-2 (PC2), respectively, which form a functional receptor-channel complex (3) that is critical for the proper regulation of cellular signaling pathways involved in kidney development and tissue repair (4).

Although the precise mechanism of cystogenesis in ADPKD remains unresolved, current evidence indicates that cyst initiation is triggered when functional polycystin levels fall below a critical threshold within the cell (5). This can result from a classical two-hit event involving a germline mutation in one *PKD* allele and a somatic loss-of-function mutation in the other allele. There is also evidence that cystogenesis can be caused by a subthreshold state due to hypomorphic variants and other genetic modifiers, or contextual factors such as developmental timing and tissue injury, any of which can lower polycystin dosage sufficiently to trigger cyst formation (1, 5). This “threshold” model has important therapeutic implications, suggesting that strategies aimed at increasing functional polycystin levels by modulating the PKD genes could substantially reduce disease progression or even halt cystogenesis. Several recent studies support the concept of targeting polycystin levels in PKD. Lakhia et al. (6) showed that increasing PC1 and PC2 protein expression by deleting miR-17 binding motifs in the 3’-untranslated regions (UTRs) of *Pkd1* and *Pkd2,* or by using an oligonucleotide inhibitor of miR-17, reduced disease severity in PKD mice models. Similarly, a single dose of base editors that corrected the “RC” mutation in *Pkd1^RC/RC^* mice, normalized functional PC1 levels and rescued the disease phenotype (7). Base editing and eukaryotic ribosomal selective glycosides that promote ribosomal read-through of premature termination codons in *PKD1* and *PKD2* restored polycystin levels and mitigated the cystic phenotype in an organoid model of ADPKD (8). Collectively, these findings provide proof-of-concept that elevating or normalizing polycystin dosage is a viable therapeutic strategy for inhibiting cystogenesis in PKD.

Building on this rationale, and recognizing that the extensive mutational heterogeneity of ADPKD (2, 9) necessitates a diverse repertoire of dosage-restoring strategies, we sought to increase PC1 levels by transcriptionally activating endogenous *PKD1* using the CRISPR activation (CRISPRa) system. CRISPRa employs a nuclease-deficient Cas9 (dCas9) tethered to transcriptional activators that, when directed to promoter or enhancer regions by guide RNAs, enable robust transcriptional activation of the target gene (10). For this study, we used dCas9 fused to the tripartite transcriptional activator VPR (VP64, p65, and Rta) and screened multiple single-guide RNAs (sgRNAs) to identify candidates with the highest potential to enhance endogenous *PKD1* expression in various cellular models of PKD. We found that the *Pkd1* CRISPRa system increased PC1 levels, even from a hypomorphic allele, and elicited beneficial effects, including the attenuation of cystogenic signaling and the cystic phenotype.

## Materials and methods

### Mammalian cell culture

HEK293T cell line (CRL-3216) was purchased from the American Type Culture Collection (ATCC) and LentiX 293T cells were purchased from Takara (632180). Both HEK293T and LentiX cells were maintained in high-glucose Dulbecco’s Modified Eagle Medium (DMEM; Sigma, D6429) supplemented with 10% fetal bovine serum (FBS; R&D Systems, S11550). M1 mouse renal cortical collecting duct cells were maintained in a 1:1 mixture of DMEM-low glucose (Sigma, D5523) and Nutrient Mixture F-12 Ham (Sigma, N6760), 15 mM HEPES, and 20 mM sodium bicarbonate and 1% penicillin-streptomycin solution (DME/F12 media), supplemented with 5% FBS. *Pkd1^RC/-^*mouse collecting duct-derived immortalized epithelial cell line (6) was a gift from Dr. Ronak Lakhia at University of Texas Southwestern Medical Center. These cells carry one null *Pkd1* allele and one hypomorphic allele (p.R3277C, abbreviated as “RC”). Mouse primary kidney tubular epithelial cells were generated from 2-week-old *Pkd1^RC/Cond^; Pkhd1^Cre-^* and *Pkd1^RC/Cond^; Pkhd1^Cre+^* mice at the PKD Rodent Model and Drug-Testing Core at KUMC. These cells were compound heterozygous for *Pkd1* carrying one “RC” allele while the other allele was either floxed (*RC/fl* control cells) or null following Cre-mediated recombination (*RC/-* cells). Primary normal human kidney (NHK) cells and human ADPKD cystic epithelial cells were obtained from the PKD Research Biomaterials and Cellular Models Core at KUMC. All primary cells and the *Pkd1^RC/-^* cells were maintained in DMEM/F12 medium supplemented with 2 × 10^-9^ M triiodothyronine (T3; Sigma, T2877), 1× insulin, transferrin and selenium premix (ITS; Corning, 354350) and 3% FBS for the *Pkd1^RC/-^* cells and 5% FBS for mouse and human primary cells. All cultures were incubated at 37°C and 5% CO_2_ (balanced with O_2_) in a humidified incubator. A summary of the various cell lines and primary cells used in this study and the purpose of each is in Supplemental Table S1.

### Guide RNA design and cloning

Mouse or human proximal *PKD1* promoter up to 500 bp upstream of the transcriptional start site (TSS) was identified in the Eukaryotic Promoter Database (11). This region was analyzed using the CRISPOR tool (12) to identify candidate sgRNA sequences containing the canonical ‘NGG’ protospacer adjacent motif (PAM). Searches were performed against the GRCm39 (mouse) or GRCh38 (human) genome assemblies (Supplemental Table S2 and S3). To refine TSS identification and account for alternative promoter usage, the promoter coordinates were validated against the FANTOM5 CAGE Promoter Atlas (13), which identified four distinct putative TSS for mouse *Pkd1*. All four TSS were incorporated into the sgRNA design strategy. To ensure a valid control, we used a custom Python script to select a non-targeting (NT) sgRNA that lacked mouse/human genomic homology and Pol III termination motifs while maintaining a GC content consistent with our *PKD1*-targeting guides. Potential off-target binding sites were calculated using Cas-OFFinder (14). The sgRNAs were cloned and expressed from the pMLM3636 (gift from Keith Joung; Addgene, 43860) and will be referred to as *Pkd1*-sgRNAx (for mouse) and *PKD1*-sgRNAx (human) where ‘x’ denotes a specific sgRNA number. Briefly, complementary oligonucleotides (from Integrated DNA Technology) encoding each 20-base protospacer sequence with 5′ phosphorylation and overhangs compatible with the pMLM3636 type IIS sites were annealed. The pMLM3636 was linearized with *Esp3I* enzyme (NEB, R0734S), dephosphorylated with shrimp alkaline phosphatase (NEB, M0371S), and gel-purified. Annealed oligos were ligated into the linearized vector and transformed into XL-1 Blue competent cells (Agilent, 200249). Positive clones were identified by Sanger sequencing (https://www.azenta.com/) across the U6-sgRNA insertion junctions.

The U6 promoter and the sgRNA sequences were also cloned into Lenti-GG-hUbc-dsRED plasmid (gift from Charles Gersbach; Addgene plasmid, 84034) using Gibson assembly (NEB, E5510S) to create individual mouse lentiviral sgRNA transfer plasmids (pLV-*Pkd1*-sgRNA-hUbc-dsRED) that were maintained in Sure 2 supercompetent cells (Agilent, 200152). All plasmids were purified using the Quantum Prep Plasmid Midiprep Kit (BioRad, 7326120) and verified by Nanopore whole-plasmid sequencing (https://www.azenta.com/) before transfection or transduction.

### Multiple sequence alignment and pairwise alignment

To evaluate the off-target risk profile of selected *PKD1*-sgRNAs, we performed pairwise sequence alignment analysis between the 12 bp seed sequences of the sgRNA and the promoter regions of the six *PKD1* pseudogenes. Local alignments were generated using EMBOSS Water (15) and the Smith-Waterman algorithm (16) to calculate percent similarity, mismatch frequency, and gap occurrence. The presence of a “NGG” PAM was assessed immediately 3’ of the aligned target sites and recorded as a binary score, where 100 indicated a functional “NGG” PAM and 0 indicated its absence.

For global promoter analysis, the 2 kb and 200 bp sequences upstream of the TSS for the functional *PKD1* and its pseudogenes were retrieved from the Ensembl database 100 (15). Multiple Sequence Alignment (MSA) was performed using Clustal Omega (15) to determine the percent identity and quantify the degree of regulatory sequence divergence across the *PKD1* paralog family.

### Lentivirus production

Lentiviral particles encoding dCas9-VPR (lenti-dCas9-VPR) or mouse *Pkd1-*sgRNAs (lenti-*Pkd1-*sgRNA) were produced using the One-Shot Lenti VSV-G system (Takara, 631275) according to the manufacturer’s instructions. Briefly, LentiX 293T cells (Takara, 632180) were transfected with the pLV-EF1α-dCas9-VPR-P2A-Puro transfer plasmid (a gift from Kristen Brennand; Addgene, 99373) or pLV-*Pkd1-*sgRNA-hUbc-dsRED together with the system’s packaging components. Culture medium was replaced 12 h post-transfection. Viral supernatants were collected at 48 h and 72 h post-transfection, pooled, passed through a 0.45 µm polyethersulfone filter, and concentrated using Lenti-X Concentrator (Takara, 631231) per the manufacturer’s protocol. Concentrated virus preparations were aliquoted and either used immediately or stored at −80°C. Viral genome titers were quantified from frozen aliquots using the Lenti-X qRT-PCR Titration Kit (Takara, 631235) following the manufacturer’s instructions.

### Generating stable HEK293T and M1 cell line constitutively expressing dCas9-VPR

HEK293T and M1 cells were transduced with lenti-dCas9-VPR at a multiplicity of infection (MOI) of 10. For spinfection, 1 × 10^6^ cells were seeded per well in 6-well plates in the presence of 8 µg/mL polybrene and the appropriate volume of concentrated lentivirus to achieve MOI 10. Plates were centrifuged at 1,000 × g for 2 h at 33°C. Following spinfection, pre-warmed culture medium was added and cells were incubated for 24 h at 37°C in a humidified incubator with 5% CO_2_. After 24 h, the medium was replaced. At 72 h post-spinfection, cells were replated into 150-mm dishes and selected with 1.5 µg/mL puromycin. Stable, single-cell-derived clones of HEK293T (HEK-dCas9-VPR) and M1 (M1-dCas9-VPR) cells constitutively expressing dCas9-VPR were isolated using the cloning cylinder method (17). The cells were maintained in their parental culture medium with 1.5 µg/mL puromycin. The expression of dCas9 was confirmed via Western blot using an anti-Cas9 antibody (see Western blot section below). To validate the functional potency of the generated clones, we performed targeted CRISPRa of *HNF4A* in HEK-dCas9-VPR cells or *Klf15* in M1-dCas9-VPR cells. Specifically, HEK-dCas9-VPR clones were transfected with an *HNF4A*-promoter targeting sgRNA (GATTGAATTAGGGGATCT) (18) using Lipofectamine LTX with PLUS Reagent (Thermo Scientific, 15338100). Conversely, M1-dCas9-VPR clones were transduced via spinfection at an MOI of 10 with lentiviral particles encoding an sgRNA targeting the *Klf15* promoter (GGGACTCTGCGGGCTTTCAG) (19). At 72 h post-treatment, cells were harvested and analyzed for their respective gene expression levels via qRT-PCR (see RNA isolation and qRT-PCR section below).

### EGFP-reporter based sgRNA screening using flowcytometry

Nucleotide sequences comprising the 5′ UTR and 1,000 bp promoter regions of mouse *Pkd1* (ENSMUSG00000032855) and human *PKD1* (ENSG00000008710) upstream of the TSS were retrieved from the Ensembl database 100 (15) and synthesized by Azenta as gene fragments with flanking restriction sites. Both inserts carried *HindIII* (NEB, R3104T) at the 5′ end; the mouse fragment harbored *KpnI* (NEB, R3142S) at the 3′ end, and the human fragment harbored *EcoRI* (NEB, R3101T) at the 3′ end. The fragments and the promoter-less pmEGFP vector (Addgene, 36409; gift from Benjamin Glick) were digested with the corresponding enzyme pairs, gel-purified, and ligated using T4 DNA ligase (NEB, M0202S) to enable directional cloning, generating the human *PKD1*-EGFP (Supplemental Fig. S1A) and mouse *Pkd1*-EGFP (Supplemental Fig. S1B) reporter plasmids.

For sgRNA screening, selected HEK-dCas9-VPR stable monoclonal cells were transfected in 12-well plates with 60.5 fmol of the appropriate *Pkd1/PKD1*-EGFP reporter and 570.1 fmol of each corresponding *Pkd1/PKD1-*sgRNA plasmid using Lipofectamine LTX with PLUS Reagent (Thermo Scientific, 15338100). Untransfected cells served as negative controls, and cells transfected with pcDNA-EGFP (Addgene, 13031; gift from Doug Golenbock) were used to define the EGFP-positive gate. Reporter-only transfected cells were included as negative controls to establish baseline autofluorescence. At 72 h post-transfection, the percentage of EGFP-positive cells (%EGFP+) was quantified by flow cytometry on an Attune NxT Flow Cytometer (Thermo Scientific). At least 10,000 singlet events were acquired per sample using Attune Cytometric Software v7.1, and data were analyzed in FlowJo v10.10. Debris was excluded using FSC-A versus SSC-A, singlets were selected using FSC-A versus FSC-H, and EGFP was measured in the singlet population on the BL1 detector (530/30 nm bandpass); the same voltages and gates were applied to all samples. Values were baseline-subtracted from the %EGFP+ and the mean fluorescence intensity (MFI) measured in reporter-only controls. The %EGFP+ and MFI for all sgRNAs were then normalized to the NT sgRNA control to calculate fold changes, which were log_2_-transformed. sgRNAs were ranked based on their EGFP activation score, calculated as log_2_ (%GFP^+^ × MFI).

### Comparative analysis of sgRNA candidates for CRISPRa of endogenous *PKD1* in immortalized cell lines

Top-performing sgRNAs from the reporter assay were evaluated for their ability to transactivate endogenous *PKD1* in HEK-dCas9-VPR or M1-dCas9-VPR cell lines. In HEK-dCas9-VPR cells, each *PKD1-*sgRNA and a NT-sgRNA were tested individually by lipofecting 2.5 µg of *PKD1-*sgRNA per well in 6 well plates. To assess mouse *Pkd1* activation, lenti-*Pkd1-*sgRNAs were used to transduce M1-dCas9-VPR cells by spinfection at an MOI of 10. At 72 h post-treatment, endogenous *PKD1* expression was quantitated via qRT-PCR using species-specific *PKD1* mRNA-primers.

### CRISPRa of endogenous *Pkd1* in mouse and human cellular PKD models

To attain sufficient transfection efficiency in cells that were difficult to transfect, we generated a miniaturized circular CRISPRa vector (mdCas9-VPR) by enzymatic excision and intramolecular ligation of the mammalian expression cassette (promoter, dCas9-VPR coding sequence, and poly(A) signal) from a full-length dCas9-VPR plasmid backbone using a method developed in our laboratory (20). Briefly, an ∼11 kb full-length dCas9-VPR plasmid was engineered with *SapI* type IIS sites flanking the expression cassette so that excision produced complementary cohesive ends on the cassette. Simultaneous T4 DNA ligase–mediated intramolecular ligation favored circularization of the excised cassette while fragmenting the vector backbone into smaller linear pieces. Linear byproducts were removed by T5 exonuclease digestion, and the circular mdCas9-VPR band was gel-purified, pooled, concentrated, quantified, and used for nucleofecting *Pkd1^RC/-^*cells, primary *RC/-* and *RC/fl* cells, NHK, and ADPKD cystic epithelial cells.

For CRISPRa of *Pkd1* in the *Pkd1^RC/-^* cell line or primary *RC/-* cells, equimolar amounts (not exceeding 136 fmol total) of *Pkd1*-sgRNAx or NT-sgRNA and mdCas9-VPR vector were nucleofected into 2.5 × 10^5^ cells per well in 24-well plates. In all conditions, total DNA mass was maintained by co-nucleofection with pBlueScript II (Agilent, 212207). Primary *RC/fl* cells were nucleofected in parallel with the mdCas9-VPR vector and NT-sgRNA and served as baseline control. Nucleofection was performed with the Ingenio Electroporation Kit (MirusBio, MIR50118) on an Amaxa Nucleofector II/2b using program T030. Cells were allowed to recover in warm complete medium, then incubated at 37°C in a 5% CO_2_ humidified incubator. Due to the difference is the recovery period between the two cell types, the *Pkd1^RC/-^* cells were analyzed at 48, 72, and 96 h post-nucleofection whereas, the mouse primary cells were analyzed at 72, 96, and 120 h post-nucleofection.

NHK and ADPKD cystic epithelial cells were nucleofected with either *PKD1-*sgRNA or NT-sgRNA using the Basic Nucleofector Kit for Primary Mammalian Epithelial Cells (Lonza, VPI-1005) on an Amaxa Nucleofector II/2b with program U-107, following the same general workflow as the murine cells. These cells were also incubated at 37°C and harvested 120 h later for qRT-PCR using *PKD1*-specific primers.

### RNA isolation and qRT-PCR

Total RNA was isolated from the cells using the Direct-zol RNA Micro Kit (Zymo, R2062) and was reverse-transcribed with the High-Capacity cDNA Reverse Transcription Kit (Applied Biosystems, 4374966). qRT-PCR was performed using SYBR Green Universal Master Mix (Applied Biosystems, 4309155) on a CFX96 Real-Time PCR Detection System (Bio-Rad). *GAPDH* served as the housekeeping gene. For human *PKD1,* the primers were designed to span distal exons 39 and 40 ensuring only the full-length *PKD1* mRNA was quantitated. Primer details can be found in Supplemental Table S4.

### Total cellular membrane preparation and immunoblotting

Total cellular membranes were prepared as described earlier (21). Briefly, nucleofected cells were seeded in 6-well plates, incubated at 37°C or 33°C in a 5% CO_2_ humidified incubator, grown to confluence, scraped from six wells and pooled. Cells were washed, resuspended in low ionic strength lysis buffer, and froze at −80°C. Frozen cells were thawed and homogenized on ice in a Dounce homogenizer. Nuclei and debris were removed by low-speed centrifugation, and membrane fractions were pelleted by high-speed centrifugation. Membrane proteins were resuspended in immunoprecipitation buffer and either stored at −80°C or processed immediately for SDS-PAGE and immunoblot analysis.

For PC1 immunoblotting, solubilized membrane protein was mixed with 4× NuPAGE LDS Sample Buffer (Invitrogen, NP0007) containing TCEP (Invitrogen, 77720), incubated at 60°C for 10 min, and separated on a NuPAGE 3–8% Tris-Acetate gel (Invitrogen, EA03785). Proteins were transferred to 0.45 µm nitrocellulose (Bio-Rad, 1620116) using the Trans-Blot Turbo RTA kit (Bio-Rad, 1704271), blocked with 2.5% fat-free milk, and probed overnight at 4°C with 7E12 (IgG1κ) anti-PC1 (gift of Dr. Christopher Ward, KUMC). Membranes were then incubated with anti-mouse IgG1 HRP (human-absorbed; Southern Biotech, 1070-05) for 1 h at room temperature and developed with a laboratory chemiluminescent substrate.

For all other targets, cells were directly lysed in RIPA buffer at 72 h post-nucleofection. Lysates were cleared by centrifugation and an aliquot equivalent to 30 µg protein was mixed with 4× NuPAGE LDS Sample Buffer containing TCEP, incubated for 5 min at 98°C, resolved on 4–15% Mini-PROTEAN TGX gels (Bio-Rad, 4561084), and transferred to 0.2 µm nitrocellulose membranes. Membranes were blocked with 5% BSA, probed overnight at 4°C with primary antibodies (1:1,000), incubated the next day with HRP-conjugated goat anti-rabbit or anti-mouse secondary antibodies (1:10,000), and developed with SuperSignal West Pico PLUS (Thermo Scientific, 34580). The following primary antibodies were purchased from Cell Signaling Technology (CST): cMyc (CST, 5605), pCreb (CST, 9198S), total-Creb (CST, 9197S), pYap1 (CST, 13008), total-Yap1 (CST, 14074), pErk (CST, 9101S), total-Erk (CST, 9102S). Other antibodies were anti-Gapdh (Santa Cruz, SC25778) and anti-Cas9 (Abcam, ab191468).

### Cell proliferation

*Pkd1^RC/-^* and primary *RC/-* cells were nucleofected with CRISPRa reagents and cell proliferation was assessed using the Cell Counting Kit-8 (CCK-8; Enzo, ALX-850-039). To ensure adequate cellular recovery post-nucleofection, cells were maintained in culture medium for 24 h prior to being detached and seeded into 96-well plates at a density of 3,000 cells/well. Cell proliferation in *Pkd1^RC/-^* cells was measured at 48, 72 and 96 h post-nucleofection. The primary cells required a more protracted recovery period and therefore were assessed at 72, 96, and 120 h post-nucleofection. At each time point, 10 µl CCK-8 reagent was added to each well and plates were incubated for 90 min at 37°C. Absorbance was read using a spectrophotometer at 450nm and the values are reported directly after background subtraction. Primary *RC/fl* cells were included as a non-PKD baseline control to compare proliferation rates with the CRISPR activated *RC/-* cells.

### 3D microcyst assay

*Pkd1^RC/-^* cells were nucleofected with CRISPRa reagents. 72 h post nucleofection, the cells were detached and suspended in ice-cold Matrigel (Corning, 354234) at 1,000 cells per 30 µL droplet, plated as hanging drops in 6-well plates, allowed to solidify at 37°C, and overlaid with defined medium containing 1:1 ratio of RenaLife basal media (Lifeline Cell Technology, LM-0010) and Advanced MEM (Thermo Scientific, 12492013), T3, 1× ITS, and 50 ug/ml Hydrocortisone (Sigma, H0888-1G). Medium was refreshed every other day for 14 days. For each experimental group, cells were plated in triplicate 30 µL hanging drops to serve as technical replicates. On day 15, droplets were fixed in 1% formalin + 1% glutaraldehyde in PBS at room temperature. Cysts within Matrigel droplets were imaged on an inverted microscope with a 2× objective using Image Pro Premier (Media Cybernetics) with extended depth-of-field and cyst areas were quantified in FIJI (22).

### MitoTracker analysis

Mitochondrial membrane potential was assessed using MitoTracker Red CMXRos (Thermo Scientific, M7512). *Pkd1^RC/-^* cells were nucleofected with CRISPRa reagents and 72 h post-nucleofection, cells were detached and reseeded into 12-well glass-bottom plates (Cellvis, P12-1.5H-N) to reach ∼70–80% confluency the next day. 24 h after reseeding, cells were incubated with 100 nM MitoTracker Red CMXRos in serum-free M1 medium for 15 min at 37°C. Immediately after staining, dye-containing medium was replaced with regular growth medium and nuclei were counterstained with 0.8 µg/mL Hoechst 33342 (Thermo Scientific, 62249) for 15 min. Cells were then washed, returned to growth medium, and imaged on a Nikon Eclipse Ti2 microscope. All images within each experiment were acquired using identical exposure settings and analyzed using NIS-elements viewer (v5.22.0). MitoTracker Red fluorescence intensity, which correlates with mitochondrial membrane potential, was quantified using FIJI (22) on five randomly selected 10X images for each replicate and used for comparison between conditions.

### Glycolytic stress test

*Pkd1^RC/-^* cells were nucleofected with CRISPRa reagents and 72 h post-nucleofection, cells were detached and reseeded into XFe96 cell culture microplates at 5000 cells per well and incubated overnight. On the day of the assay, the growth medium was replaced with warmed glucose-free Seahorse XF Base Medium (Agilent, 103335-100) supplemented with 2 mM glutamine and 1 mM sodium pyruvate. Cells were incubated in a non-CO_2_ incubator at 37°C for 1 h prior to the assay to achieve baseline glucose starvation. Extracellular Acidification Rate (ECAR) was measured using XF Glycolysis Stress Test Kit (Agilent) on an XFe96 Analyzer (Agilent) following the manufacturer’s instructions. Following the assay, cells were counterstained using Hoechst 33342 dye, and total cell counts per well were quantified on a Cytation cell imaging multi-mode reader to normalize acidification rates per 1,000 cells. The following parameters were calculated using the maximum rate achieved within each measurement cycle.

Basal Glycolytic Rate = ECAR after glucose − ECAR before glucose

Maximum Glycolytic Capacity = ECAR after oligomycin − ECAR before oligomycin

Glycolytic Reserve = Maximum Glycolytic Capacity − Basal Glycolytic Rate

### Cyclic AMP assay

Intracellular cyclic AMP (cAMP) levels were quantified using a competitive ELISA kit (Enzo Life Sciences, ADI-900-066A). 72 h after nucleofection, cells were lysed in 0.1M HCl and 0.5% Triton X-100 to inhibit endogenous phosphodiesterase activity, and supernatants were acetylated with acetic anhydride and triethylamine before processing them according to the manufacturer’s instructions. Final cAMP concentrations were calculated using a 4-parameter logistic (4PL) curve fit and normalized to the total cellular protein content of each sample.

### Statistics

Statistical analyses were performed, and figures were generated using GraphPad Prism v9 (GraphPad Software, LLC). For qRT-PCR data, statistical analysis was performed on -ΔCq. In experiments comparing multiple sgRNAs within a cell type, significance was determined by repeated measures one-way ANOVA followed by Dunnett’s or Tukey’s post-hoc test. In experiments involving comparisons across different cell types or genotypes (e.g., Fig. 5A), an ordinary one-way ANOVA followed by Tukey’s multiple comparisons test was employed. For time-course experiments involving multiple variables (e.g., CCK-8 assays), a two-way ANOVA followed by Sidak’s or Tukey’s post-hoc test was used. For comparisons between two groups, a two-tailed paired t-test was used when data were aggregated per biological replicate. For hierarchical data involving multiple measurements within each biological unit (e.g., individual cyst areas), a nested t-test was used to account for technical replicates within biological replicates. All experiments were performed with at least three independent biological replicates. A p value of less than 0.05 was considered statistically significant. Specific statistical tests for supplemental data are indicated in the respective figure legends.

## Results

### Generation of a CRISPRa platform for sgRNA screening

We generated and validated a HEK-dCas9-VPR stable monoclonal cell line to establish a robust CRISPRa platform for screening sgRNAs (Supplemental Fig. S2A). Functional validation with an *HNF4A* targeting sgRNA (18) identified clone 4 as having the highest transactivation potential (Supplemental Fig. S2E). This clone was therefore selected for all subsequent *PKD1-*sgRNA screening experiments.

We designed 11 human *PKD1*-sgRNAs (Fig. 1A; Supplemental Table S2) and 15 mouse *Pkd1*-sgRNAs (Fig. 1A; Supplemental Table S3). These were screened by co-transfecting them together with their respective human or mouse *PKD1* promoter-EGFP reporter into the validated HEK-dCas9-VPR cell line (Fig. 1B). The performance of each sgRNA was evaluated via flow cytometry and ranked according to their EGFP activation scores. All *PKD1*-sgRNAs demonstrated robust EGFP activation (10^3^- to 10^4^-fold increase over NT-sgRNA), from which the four top-performing guides (sgRNAs 10, 12, 15, and 18) were selected for downstream validation (Fig. 1C and E). Similarly, the five highest-ranking mouse *Pkd1* guides (sgRNAs 11–15) exhibited a consistent ∼10^3^-fold increase in EGFP activation over the NT-sgRNA and were prioritized for further characterization (Fig. 1D and F).

**Fig. 1.**
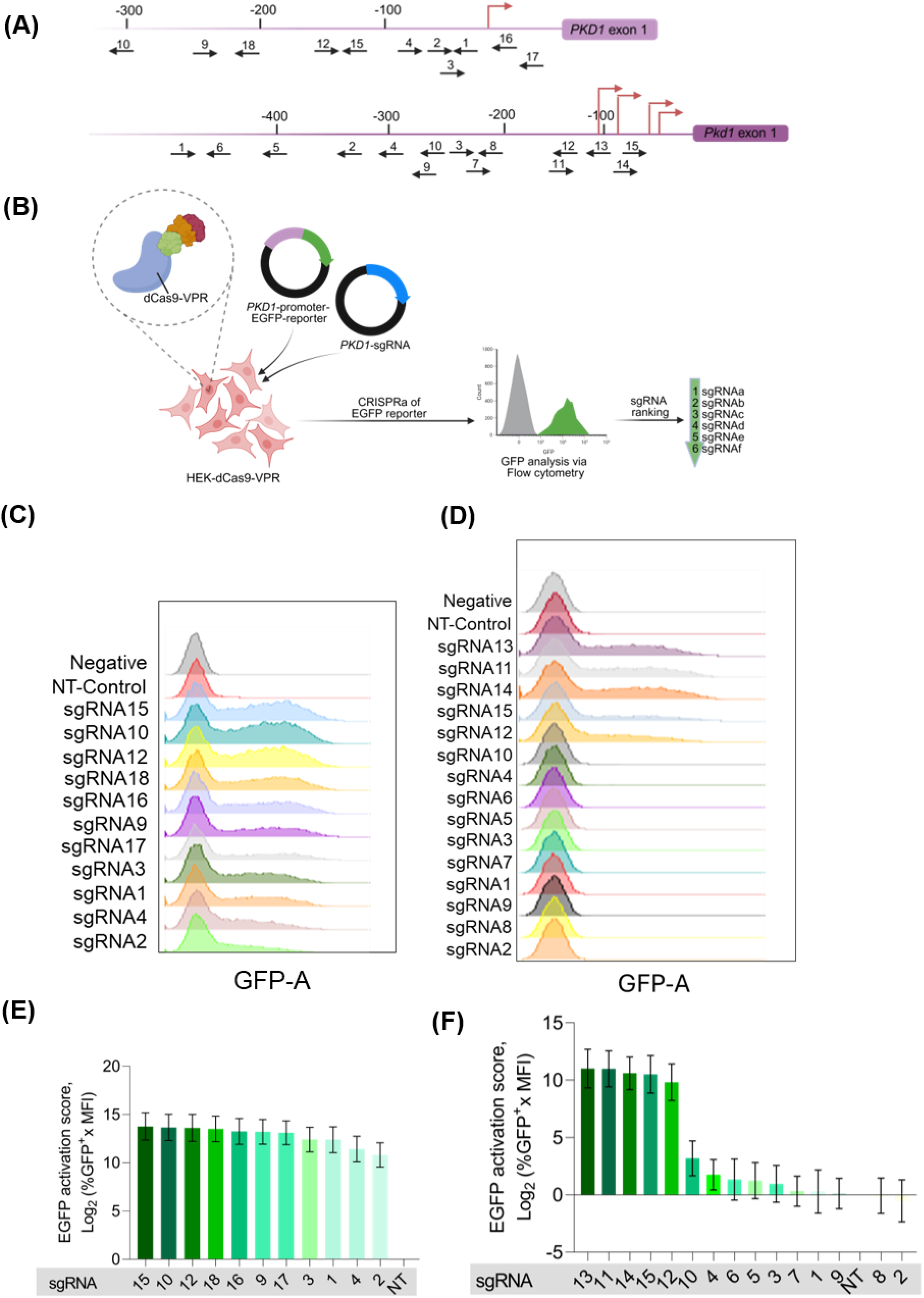
Screening of potential guide RNA sequences for CRISPRa of human and mouse *PKD1*. **(A)** Schematic representation of human *PKD1* (top) and mouse *Pkd1* (bottom) relative to the transcription start site (TSS, red bent arrow). For the mouse *Pkd1* promoter, four putative TSSs are shown; genomic coordinates (–100 to –400 bp) are relative to the TSS most proximal to exon 1. Numbered black arrows indicate the positions of sgRNA target sites; arrow direction denotes strand orientation (right-facing: sense; left-facing: antisense). **(B)** Schematic of the experimental workflow for identifying optimal mouse or human *PKD1*-targeting sgRNAs in HEK-dCas9-VPR stable cells using mouse or human *PKD1*-promoter-driven EGFP reporter plasmid and individual sgRNA expression vectors. EGFP expression was quantified by flow cytometry and sgRNAs were ranked based on their ability to activate the EGFP reporter. **(C–D)** Representative flow cytometry histograms of EGFP fluorescence for **(C)** human and **(D)** mouse sgRNAs, ordered by decreasing EGFP activation score except NT-control sgRNA that is shown for reference. **(E–F)** EGFP activation scores (Log₂ [%GFP⁺ × MFI]) for all **(E)** 11 human *PKD1*-sgRNAs and **(F)** 15 mouse *Pkd1*-sgRNAs, ranked in descending order. Data represents mean ± SD (N = 3 independent biological replicates). NT= Non-targeting, EGFP= Enhanced green fluorescent protein, MFI= Mean fluorescence intensity

### Selection of optimal sgRNAs for endogenous *PKD1* activation

To assess the CRISPRa efficiency of the top-performing human *PKD1*-sgRNAs in activating endogenous *PKD1*, the four lead sgRNAs identified from the reporter screen (sgRNAs 10, 12, 15, and 18) were individually transfected into HEK-dCas9-VPR stable cells. Endogenous *PKD1* mRNA expression quantified by qRT-PCR, revealed that two of the four sgRNAs significantly upregulated *PKD1* relative to the NT-sgRNA, with activation levels ranging from 2.75- to 4-fold (Fig. 2A). Based on these results, sgRNA12 and sgRNA15 were prioritized for further experiments.

**Fig. 2.**
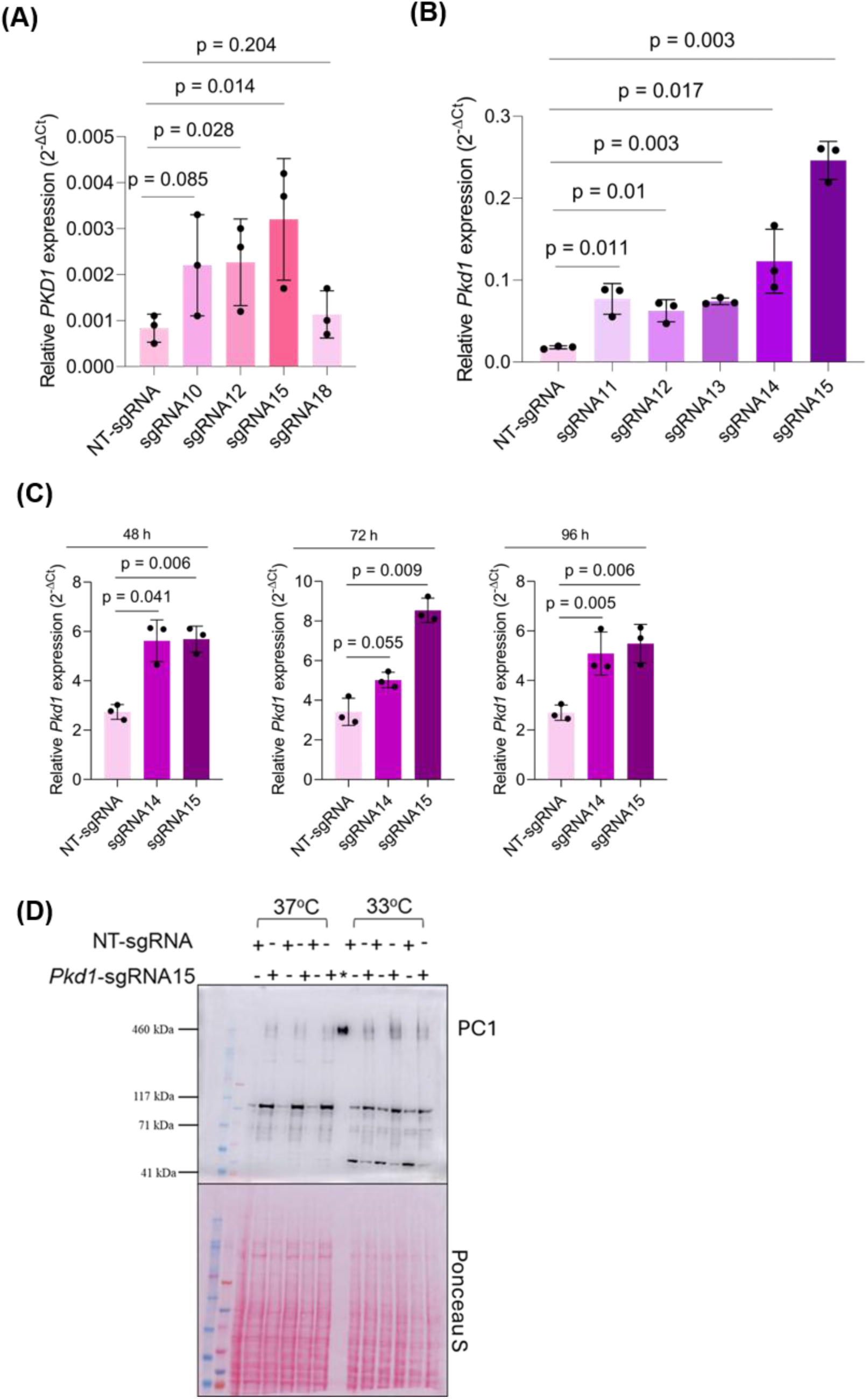
Validation of selected guide RNAs for *PKD1* CRISPRa in human and mouse cell lines. **(A-B)** Relative *PKD1* mRNA expression in **(A)** HEK-dCas9-VPR stable cells and **(B)** M1-dCas9-VPR stable cells following treatment with either NT-sgRNA or indicated sgRNAs. **(C)** Relative *Pkd1* transcript levels in *Pkd1^RC/-^* cells following nucleofection with mdCas9-VPR and indicated sgRNAs at 48, 72, and 96 h. For **(A-C)** data are presented as mean 2^-ΔCq^ ± SD (N=3 biological replicates). **(D)** Immunoblot of total membrane fractions from *Pkd1^RC/-^* cells incubated at 37°C or 33°C following treatment with NT-sgRNA or *Pkd1*-sgRNA15. PC1 (460 kDa) was detected using the 7E12 antibody. mRNA= messenger RNA, *Urinary exosome as positive control for PC1.

We also generated a mouse M1 cell line constitutively expressing dCas9-VPR (M1-dCas9-VPR; Supplemental Fig. S3A) to study the CRISPRa efficiency of the top-performing *Pkd1*-sgRNAs in activating endogenous *Pkd1.* Functional validation with a *Klf15* targeting sgRNA (19) identified clone 8 as having the highest transactivation potential (Supplemental Fig. S3I). This clone was therefore selected for subsequent endogenous *Pkd1* activation studies. M1-dCas9-VPR stable cells were transduced with lentiviral particles encoding the five highest-ranking mouse *Pkd1*-guides (sgRNAs 11–15). Consistent with our human cell data, all five sgRNAs significantly upregulated endogenous *Pkd1* compared to the NT-sgRNA at 72 h post-transduction, with activation levels ranging from 3.44- to 13.66-fold (Fig. 2B). Due to its superior fold induction, sgRNA15 was selected for all primary downstream experiments, while sgRNA14 was retained as a secondary candidate to ensure experimental rigor and reproducibility.

### CRISPRa of *Pkd1* in a hypomorphic mouse PKD cell model

To extend our findings to PKD-relevant cell model, we investigated whether CRISPRa could upregulate *Pkd1* expression in cells harboring a single hypomorphic *Pkd1* allele (*Pkd1^RC/-^)*. The p.R3277C (RC) mutation encodes a temperature-sensitive folding mutant of PC1 that causes improper protein folding, resulting in a ∼60% reduction in mature PC1 trafficking to the plasma membrane and primary cilia (23). However, the fraction of PC1 that successfully traffics remains functional, and recent studies have demonstrated that boosting functional PC1 expression is sufficient to rescue PKD phenotypes and attenuate cystic disease *in vivo* (6). We nucleofected *Pkd1^RC/-^* collecting duct cells with mdCas9-VPR and either *Pkd1-*sgRNA14*, Pkd1*-sgRNA15 or NT-sgRNA and quantified endogenous *Pkd1* mRNA expression at 48, 72 and 96h. At all three time points, *Pkd1*-sgRNA15 significantly upregulated *Pkd1* levels over NT-sgRNA by 2-2.5-fold (Fig. 2C). *Pkd1-*sgRNA14 also showed significant increase in *Pkd1* expression over NT-sgRNA at 48 and 96 h (∼1.8 - 2-fold increase; Fig. 2C); however, at 72h, the response did not reach significance despite the increase in expression (∼1.47-fold increase; Fig. 2C). We evaluated potential off-target transcriptional activation and “bystander” effects on neighboring genes to ensure the observed CRISPRa was specific to the *Pkd1* locus. Cas-OFFinder predicted no off-targets with <3 mismatches for sgRNA15 but two notable off-target candidates for *Pkd1*-sgRNA14: a 2-nucleotide mismatch site in the *Cds1* promoter and a 3-nucleotide mismatch site in exon 1 of *Sart3*. Analysis of *Pkd1^RC/-^* cells at 72 h post-nucleofection revealed no significant induction of *Sart3* or *Cds1* compared to NT-sgRNA controls (Supplemental Fig. S4A and B). We also investigated potential “bystander” activation of *Rab26*, whose promoter lies in close genomic proximity to the *Pkd1* locus. Despite the potency of the VPR trans-activator, *Rab26* expression remained unaltered following nucleofection with either *Pkd1-*sgRNA14 or *Pkd1-*sgRNA15 (Supplemental Fig. S4C and D).

To determine whether the transcriptional activation of *Pkd1* could induce endogenous PC1 protein expression from the “RC” allele, total cellular membrane fractions were prepared for immunoblot analysis. As the p.R3277C (“RC”) variant is a temperature-sensitive folding mutant that exhibits enhanced trafficking at 33°C compared to 37°C (23), we evaluated expression at both temperature. A band was observed at ∼460 kDa corresponding to full-length PC1 in cells expressing *Pkd1*-sgRNA15, whereas no PC1 signal was detected in NT-sgRNA controls at both 33°C and 37°C (Fig. 2D).

### Effects of *Pkd1* CRISPRa on disease-associated signaling, cell proliferation and cyst formation

We investigated whether the upregulation of endogenous *Pkd1* could reverse several well-known pathogenic events associated with PKD using several independent assays. We first examined the effect of *Pkd1* CRISPRa on the proliferative phenotype of PKD cells using a CCK-8 assay. While cell densities remained comparable at early time points, *Pkd1*-sgRNA15-treated cells exhibited a significant attenuation in proliferation compared to NT-sgRNA controls at both 72 h and 96 h post-nucleofection (p < 0.05; Fig. 3A). This anti-proliferative effect translated to a profound reduction in cystogenic potential in a 3D microcyst assay. *Pkd1*-sgRNA15 treatment resulted in a significantly lower total number of cysts compared to NT-sgRNA controls (p = 0.039; Fig. 3B and C), although the mean surface area of the remaining cysts did not differ between groups (Fig. 3D). We further investigated whether these phenotypic improvements were accompanied by a restoration of mitochondrial health. Quantitative fluorescence microscopy revealed that *Pkd1* CRISPRa significantly improved mitochondrial membrane potential, as evidenced by an increase in MitoTracker Red CMXRos mean fluorescence intensity (MFI) (p = 0.026; Fig. 3E and F).

**Fig. 3.**
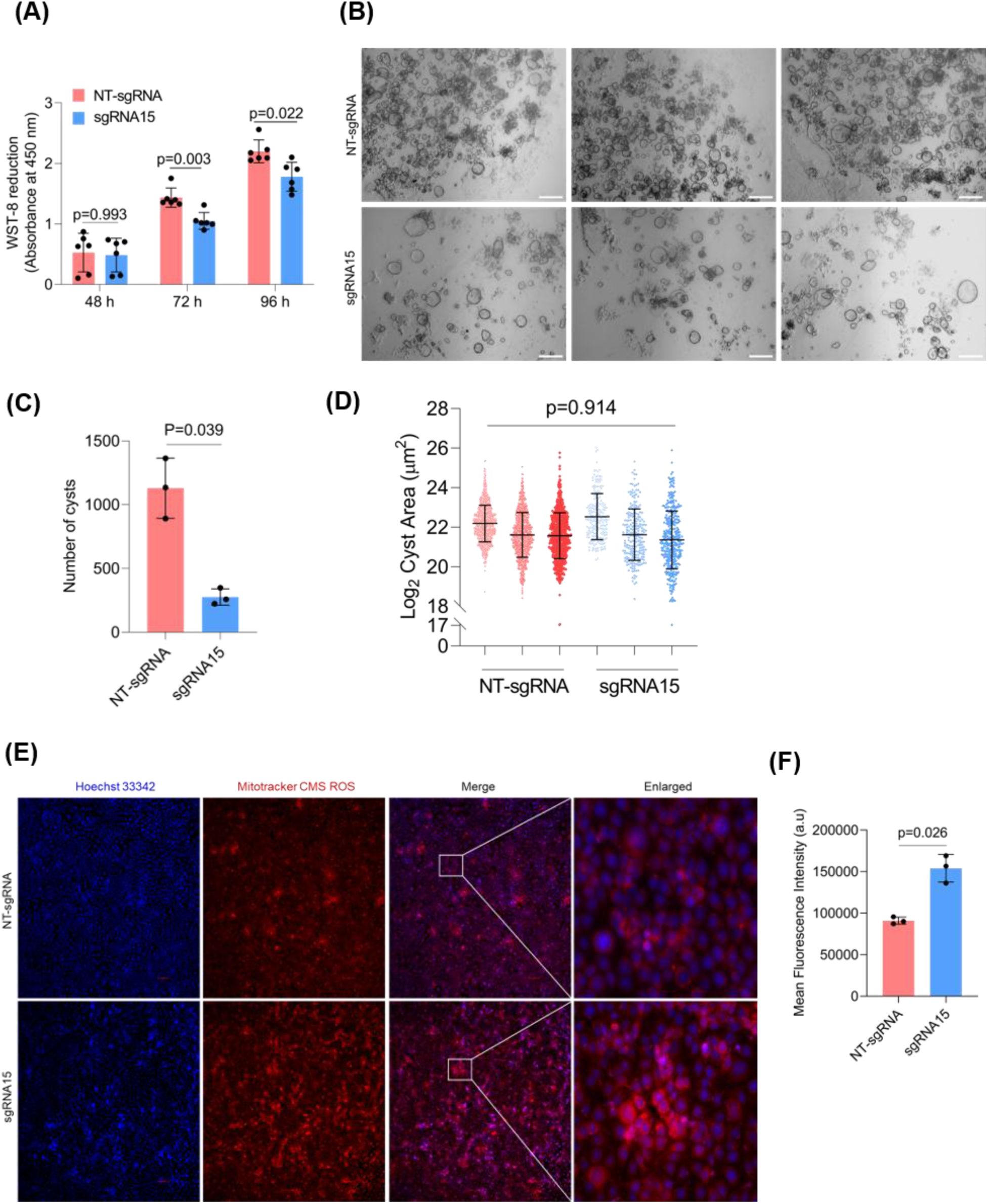
Effect of *Pkd1* CRISPRa on disease phenotypes in mouse PKD cells. **(A)** Growth kinetics in *Pkd1^RC/-^*cells were assessed at 48, 72, and 96 h post-nucleofection via CCK-8 assay (N=6 biological replicates). Data represent mean absorbance ± SD. **(B)** Representative phase-contrast images of 3D microcysts. *Pkd1^RC/-^* cells were seeded in Matrigel 72 h post-nucleofection, and cyst growth evaluated after 14 days. Images (2X magnification) show representative quadrants from N=3 independent biological replicates, with NT-sgRNA (top) and *Pkd1*-sgRNA15 treated cells (bottom). Scale bar = 1000 µm. **(C)** Quantification of total number of microcysts per experiment. Each data point represents the aggregate count from three technical replicate droplets. **(D)** Individual cyst cross-sectional area (µm^2^) on a log_2_ scale and grouped by treatment and experiment. **(E, F)** Representative images **(E)** and quantification **(F)** of MitoTracker Red CMXRos fluorescence intensity 96 h post-nucleofection with mdCas9-VPR and either NT-sgRNA or *Pkd1*-sgRNA15. Mean fluorescence intensity (MFI) was calculated from five randomly selected 10X fields per replicate and marked areas are enlarged in adjacent panels. For **(C, D and F)** data are presented as mean ± SD (N= 3 independent biological replicates).

Enhanced glycolysis is a common feature of the Warburg phenotype that is exhibited by PKD cells because of metabolic reprogramming (24, 25). We therefore investigated the effect of *Pkd1* upregulation on the *Pkd1^RC/-^* cells’ ability of aerobic glycolysis. Compared to NT-sgRNA controls, cells treated with *Pkd1*-sgRNA15 showed an overall attenuation of this metabolic state, marked by significant decreases in baseline glycolysis (p < 0.02) and maximum glycolytic capacity (p < 0.013; Fig. 4A and B). Here, basal glycolysis represents the rate of glycolysis in unstressed cells with a plentiful supply of glucose, maximum glycolytic capacity indicates the maximum glycolytic output achievable when oxidative phosphorylation is blocked by oligomycin, and glycolytic reserve is the difference between basal and maximum glycolytic rates. Both human and murine PKD models also exhibit elevated intracellular cAMP levels, a key driver of renal cyst growth (26–28). We found that the activation of endogenous *Pkd1* significantly reduced intracellular cAMP concentrations (p = 0.03; Fig. 4C). In addition, immunoblot analysis revealed that, in comparison to NT-sgRNA controls, *Pkd1*-sgRNA15 treatment significantly reduced levels of cMyc, phosphorylated Creb (pCreb), and phosphorylated Erk (pErk), while increasing phosphorylated Yap1 (pYap1) levels, which effectively corrected key signaling components that are dysregulated in PKD kidneys (6, 28, 29) (Fig. 4D). Collectively, these data demonstrate that the upregulation of endogenous *Pkd1* corrects cellular signaling and attenuates the cystogenic potential of PKD cells.

**Fig. 4.**
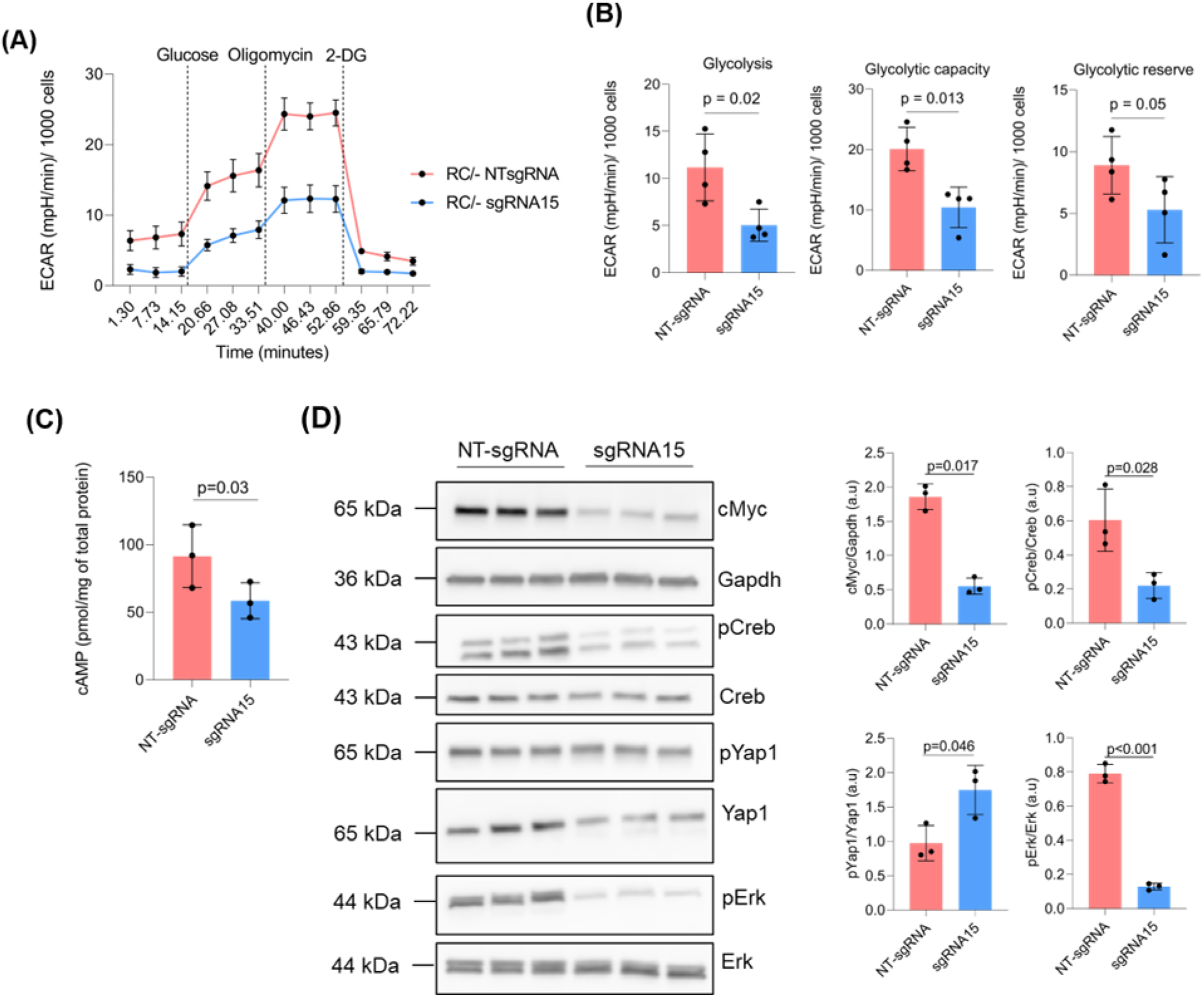
Effect of *Pkd1* CRISPRa on cellular energetics and dysregulated signaling pathways in mouse PKD cells. **(A)** Real-time extracellular acidification rate (ECAR) profile during a Seahorse XF Glycolysis Stress Test following sequential injections of glucose, oligomycin, and 2-deoxyglucose (2-DG). **(B)** Quantification of glycolytic parameters including baseline glycolysis, glycolytic capacity and glycolytic reserve. **(A-B)** Values are normalized to cell count using a scale factor of 1000 and represent mean ± SD (N=4 biological replicates). **(C)** Total intracellular cAMP quantified by ELISA in *Pkd1^RC/-^* cells. Data were normalized to total protein and are presented as mean cAMP (pmol/mg total protein) ± SD. **(D)** Representative immunoblots and densitometric quantification of cMyc, pCreb, pErk, and pYap1 expression in *Pkd1^RC/-^* cells. Protein levels were normalized to Gapdh and are expressed as mean ± SD (N=3 biological replicates).

### CRISPRa of *PKD1* in primary cellular models of PKD

To further validate our CRISPRa strategy, we determined whether our sgRNAs could activate *Pkd1* in primary PKD cells. We observed that mouse primary *RC/-* cells exhibited a 32% to 61.54% deficit in *Pkd1* expression compared to *RC/fl* cells over 72 to 120 h (Fig. 5A). Treatment of *RC/-*cells with *Pkd1*-sgRNA15 resulted in a 2.95-fold, 3.02-fold, and 3.1-fold increase in *Pkd1* transcripts relative to NT-sgRNA (p < 0.001; Fig. 5A) at 72, 96 and 120 h respectively. This robust activation significantly surpassed the levels of *Pkd1* in *RC/fl* control at 72 and 96 h post-nucleofection. *Pkd1-*sgRNA14 also induced significant activation in *RC/-* cells, increasing *Pkd1* levels over ∼2-fold relative to NT-sgRNA at all three time-points (p < 0.001; Fig. 5A) and to levels comparable to *RC/fl* control cells. We also confirmed the specificity of *Pkd1* CRISPRa in these primary cells, as evidenced by the lack of significant induction of *Sart3*, *Cds1,* and *Rab26* at 72 h post-nucleofection (Supplemental Fig. S5A-D).

**Fig. 5.**
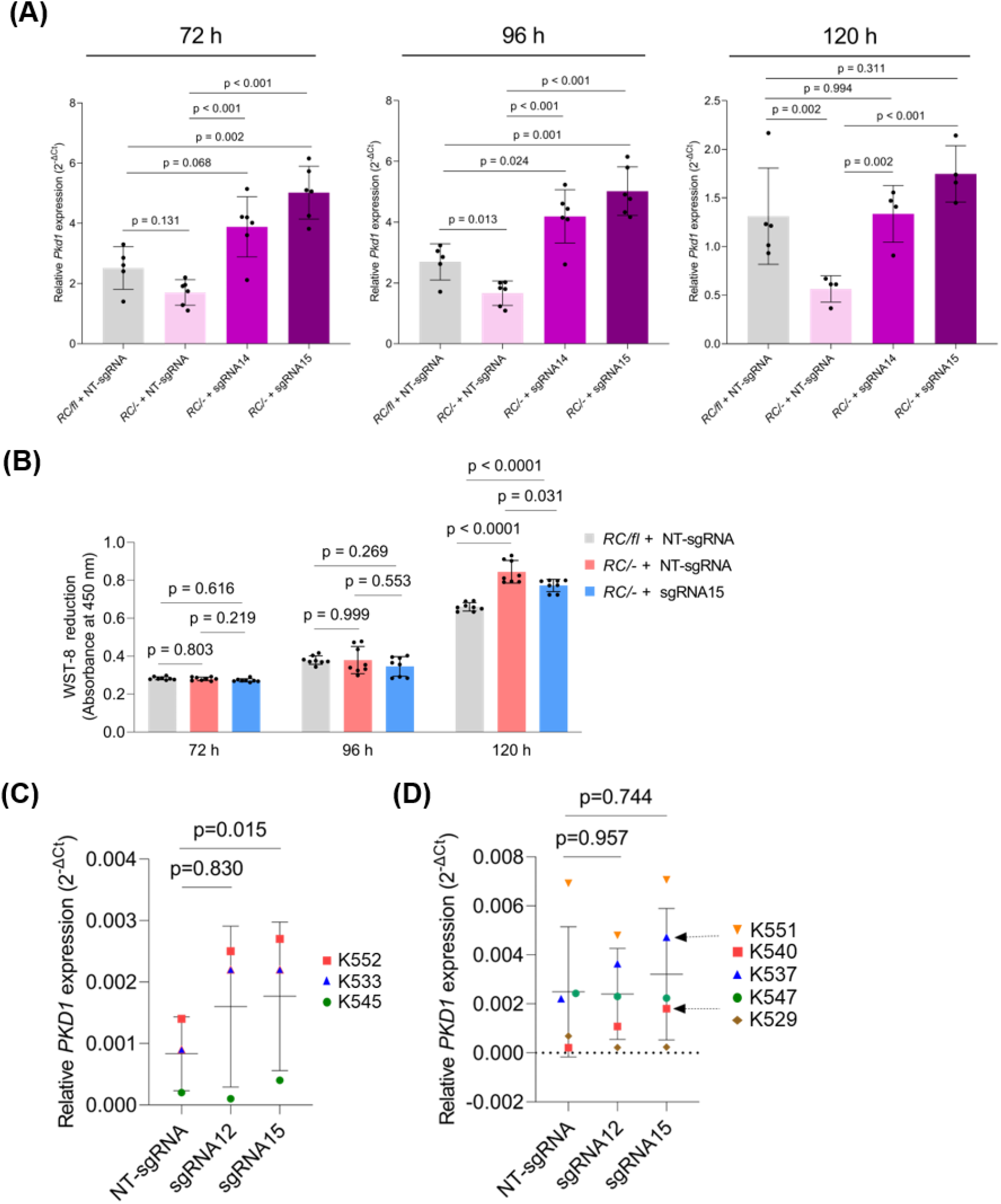
Validation of *PKD1* CRISPRa in primary mouse and human PKD cells. **(A)** Relative *Pkd1* transcript levels in primary *RC/-* cells (*Pkd1^RC/Cond^; Pkhd1^Cre+^)* following nucleofection with mdCas9-VPR and indicated mouse sgRNAs at 72, 96 and 120 h. Primary *RC/fl* cells *(Pkd1^RC/Cond^; Pkhd1^Cre-^*) nucleofected with mdCas9-VPR and NT-sgRNA served as non-PKD cell control. Data are presented as mean 2^-ΔCq^ ± SD (N = 6 for 72 and 96 h and N = 4 for 120 h). **(B)** Cell proliferation kinetics were assessed at 72, 96, and 120 h post-nucleofection in mouse primary cells using a CCK-8 assay. Data are presented as mean raw absorbance values ± SD (N = 6). **(C-D)** Relative *PKD1* transcript levels in **(C)** primary NHKs (N = 3) and **(D)** primary ADPKD cystic epithelial cells (N = 5) following nucleofection with mdCas9-VPR and indicated sgRNAs. Individual donors are represented by unique symbol and color combinations and their corresponding donor IDs as indicated. Data are presented as mean 2^-ΔCq^ ± SD. Black arrows in **(D)** indicate two “responder” lines exhibiting robust activation.

We evaluated whether *Pkd1* induction in these primary cells translated into a functional rescue of the proliferative phenotype observed in PKD cells. Cell proliferation was monitored at 72, 96, and 120 h post-nucleofection. No significant differences in cell density were observed at 72 h or 96 h. However, by 120 h the *RC/-* cells treated with NT-sgRNA exhibited a significantly higher proliferative rate compared to the *RC/fl* cells (p < 0.0001; Fig. 5B). CRISPRa of *Pkd1* with *Pkd1-*sgRNA15 in *RC/-* cells significantly attenuated this difference (p = 0.031 v NT-sgRNA).

We next tested whether our human *PKD1-*specific guides could activate endogenous *PKD1* in primary human renal epithelial cells. Primary cells harvested from normal human kidney (NHK) and cyst-lining epithelial cells from human ADPKD kidneys were nucleofected with mdCas9-VPR and either *PKD1*-sgRNA12, *PKD1*-sgRNA15, or NT-sgRNA. In NHK cells, *PKD1*-sgRNA15 induced a ∼2.1-fold upregulation of endogenous *PKD1* (p = 0.015; Fig. 5C). *PKD1*-sgRNA12 showed a non-significant ∼1.9-fold increase over NT-sgRNA. In ADPKD cyst cells derived from five independent donors, the mean increase in *PKD1* expression across all primary human donor lines did not reach statistical significance. Cells from two donors exhibited robust activation (black arrows; Fig. 4D), with sgRNA15 increasing *PKD1* mRNA by 2.1- and 8.5-fold, and sgRNA12 yielding 1.6- and 5.1-fold inductions, respectively. In contrast, the remaining three donor lines showed no significant induction with either guide RNA, highlighting the variability in CRISPRa efficiency across different primary human samples

Finally, analysis for potential bystander activation of the neighboring *RAB26* locus showed that *RAB26* expression was not significantly altered by either sgRNA compared to NT-sgRNA controls at 120 h post nucleofection (Supplemental Fig. S6A).

## Discussion

In this study, we strengthen the argument that augmenting *PKD1* expression in a PC1-deficient environment represents a viable therapeutic strategy for ADPKD. We demonstrate that CRISPRa of endogenous *Pkd1* increases PC1 levels and suppresses cystogenic signaling in mouse PKD cell models. These molecular improvements translate to significantly reduced cell proliferation and cyst formation *in vitro*. Our findings establish a compelling proof-of-concept for the use of CRISPRa to achieve targeted, endogenous *PKD1* upregulation.

While the massive size of the *PKD1* gene (∼52 kb) and its 14.5 kb transcript have historically precluded traditional cDNA-based gene replacement, recent evidence (6–8) suggests targeting the endogenous *Pkd1* locus can successfully bypass these structural hurdles. To develop CRISPRa reagents capable of tackling the root cause of ADPKD, we systematically screened and identified sgRNAs for *PKD1* transactivation. We targeted the proximal promoter within a 500 bp window upstream of the TSS, a region generally considered optimal for CRISPRa (30). However, this region in both mouse and human *PKD1* exhibits an exceptionally high GC content, exceeding 70% in mice and 90% in humans (Supplemental Fig. S6B). Such high GC density is known to detrimentally affect sgRNA binding due to the formation of complex secondary structure (30, 31). Despite these biochemical impediments, we successfully established a narrow, optimal window for potent transcriptional activation of *PKD1* in both species.

With these systematically scrutinized sgRNAs, we demonstrated that CRISPRa can effectively target and enhance transcription from a hypomorphic allele. By expanding the pool of functional PC1 protein this approach likely pushes polycystin levels beyond the critical disease threshold, leading to a significant rescue of multiple ADPKD-associated pathogenic markers. Notably, the 2.5-fold induction of *Pkd1* transcripts observed in *Pkd1^RC/-^*cells is consistent with other therapeutic CRISPRa studies, where similar magnitudes of transcriptional activation have proven sufficient to yield substantial functional benefits (32–34). The molecular consequences of this *Pkd1* transactivation were profound. The observed decrease in cystogenic stimuli, such as reduced intracellular cAMP, coupled with favorable shifts in pro-proliferative and oncogenic pathways like Creb, Erk, cMyc and Yap1 suggests that PC1 could indeed serve as an upstream master regulator of these diverse cellular cascades. On a metabolic level, while our analyses showed reduction of the glycolytic ‘Warburg’ phenotype and a restoration of mitochondrial membrane potential, these end points capture only a portion of the metabolic network. Future studies utilizing real-time oxygen consumption rate (OCR) measurements will be valuable to provide a fully integrated view of how endogenous *Pkd1* upregulation fixes mitochondrial oxidative respiration alongside glycolytic pathways. We also acknowledge that PC1 must be properly trafficked to the primary cilium to execute its essential roles in mechanosensation and ciliary signaling. While our current total membrane preparations confirm successful protein production after CRISPRa, they do not distinguish between ciliary and extraciliary PC1 pools. Therefore, future studies utilizing high-resolution imaging will be valuable to formally characterize the spatial distribution of CRISPRa-induced PC1.

Our finding that CRISPRa of the hypomorphic *RC* allele can suppress cyst formation aligns with reports showing the attenuation of cystic phenotypes upon *Pkd1* de-repression (6), mutational correction (7, 35) or pre-mature stop codon read-through (8), all of which increase the functional pool of PC1 in affected cells. Interestingly, while the total frequency of cyst initiation was dramatically reduced in the *Pkd1-*CRISPR-activated group, the remaining cysts reached mean sizes comparable to those in the NT-sgRNA controls. This observation likely reflects two distinct phenomena. First, we speculate that these cysts represent a subpopulation of cells where *Pkd1* induction did not reach the critical threshold required to inhibit cystogenesis, likely due to the inherent stochasticity of CRISPRa delivery. Second, the subsequent expansion of these cysts was likely facilitated by the significantly lower cyst density in the CRISPR-activated cultures. In this environment, reduced competition for nutrients and physical space likely permitted these cysts to reach their maximal growth potential, whereas expansion in the control cultures was limited by the nutrient-depleted, crowded environment of the high-density Matrigel droplets. Ultimately, in this experimental context, our results suggest that *Pkd1*-CRISPRa may act as a potent gatekeeper of cyst initiation, effectively preventing the initial cellular transformation to a cystic phenotype. This points to a therapeutic model where a modest prophylactic elevation of PC1 in phenotypically normal tubule cells, including those harboring germline mutations, maintains levels above the threshold required to resist cystogenesis, especially during periods of kidney injury. Future studies utilizing inducible dCas9-VPR systems to upregulate *Pkd1* in pre-established microcysts will be necessary to clarify whether CRISPRa can actively remodel existing cystic architecture, similar to the rapid reversal demonstrated by Dong et al. following genetic re-expression (36).

Expanding our proof-of-concept to primary mouse and human cells confirmed the efficacy of our CRISPRa platform. For instance, CRISPRa of *Pkd1* in mouse primary *RC/-* cells led to an attenuation of the proliferative phenotype. It is worth noting that these primary cells represent a heterogeneous population of tubular epithelial cells where complete deletion of the second *Pkd1* allele occurs only within the *Pkhd1*-driven, Cre-expressing collecting duct lineage. While future studies utilizing sorted or more homogenous populations of primary *Pkd ^RC/-^* cells would provide additional clarity, the consistent attenuation of the proliferative phenotype observed across both the tightly controlled, clonal cell line (*Pkd1^RC/-^*) and this heterogeneous primary cell system reinforces the robustness and translational potential of our CRISPRa platform. To evaluate the human translational relevance of these findings, our subsequent validation in primary NHK and ADPKD cystic cells confirmed the efficacy of our CRISPRa platform in a human genomic context. A unique challenge in activating *PKD1* is the presence of six highly homologous pseudogenes (*PKD1P1–P6*), which could potentially sequester the dCas9-activator complex. However, sequence analysis reveals significant divergence between *PKD1* and five of these pseudogenes (*PKD1P2–P6*), characterized by high mismatch rates (up to 25%) and a lack of matching PAM sequences (Supplemental Fig. S7A and B). This divergence in sequence from *PKD1* is evident throughout the 2 kB region upstream of the TSS (Supplemental Fig. S7C) and becomes particularly pronounced within the 200 bp proximal promoter (the optimal window for our CRISPRa sgRNA to bind; Supplemental Fig. S7D). While the 100% match with *PKD1P1* may contribute to some degree of dCas9 sequestration, its activation is unlikely to be pathogenic as it lacks protein-coding capacity.

It is also important to acknowledge that *PKD1* dosage is a critical safety factor, as mice expressing PC1 significantly above wild-type levels (2- to −15-fold) can, paradoxically, develop cystic disease (37, 38). Therefore, we purposefully avoided CRISPRa with pooled sgRNAs, as this has been shown to increase transcription even more than single guides (30, 39–41). Also, having a single guide not only is more cost-and labor-effective but also enhances the feasibility of viral packaging and delivery to complex organs like the kidney. It remains to be seen, however, if the robust activation observed in our cell models can be replicated *in vivo*, where factors such as epigenetic silencing, chromatin accessibility, and delivery efficiency may dampen CRISPRa performance. Future studies should explore how to best leverage the inherent “tunability” of our CRISPRa platform. By testing different sgRNA configurations, modulating the dCas9 dose, or employing transactivation domains with varying potencies (42, 43), we can determine how to reliably maintain *PKD1* expression within a safe, restorative therapeutic window. Additionally, while we performed off-target analysis on a small subset of genes, we acknowledge that this does not completely rule out broader, genome-wide transcriptional effects driven by the potent VPR activator. Therefore, future translational investigations must prioritize comprehensive, unbiased safety profiling. Utilizing genome-wide transcriptomic approaches such as RNA-sequencing (RNA-seq), combined with epigenomic mapping assays like chromatin immunoprecipitation-sequencing (ChIP-seq) or CUT&RUN, will be essential to systematically characterize the global off-target profile and ensure strict locus specificity.

Beyond these structural considerations, the evaluation of the primary human cells provided critical insights into the translational feasibility of our CRISPRa platform. We observed a heterogeneous response across both NHK and ADPKD donor lines. This variability likely reflects the biological complexity of primary human samples compared to immortalized cell lines. In instances where activation was absent, technical factors such as primary cell health, varying nucleofection efficiencies, or differences in chromatin accessibility at the *PKD1* locus may have limited dCas9-VPR activity. Specifically, for the ADPKD cohort, the underlying genotype may also dictate therapeutic success. The efficacy of CRISPRa may be contingent on the presence of a targetable *PKD1* allele, such as one harboring a hypomorphic or non-truncating mutation with some residual function, rather than large genomic deletions or promoter-disrupting rearrangements. From a clinical translation perspective, this heterogeneity indicates that clinical application of *PKD1* CRISPRa will likely require rigorous pre-treatment patient stratification. Genetic screening will be essential to identify individuals with intact, targetable promoter architectures and responsive mutant or fully functional alleles, ensuring that therapeutic interventions are directed toward patients with the highest probability of molecular rescue. Furthermore, understanding whether epigenetic variations contribute to non-responsiveness will be critical for considering transient co-therapies that enhance locus accessibility for CRISPRa.

In conclusion, our work demonstrates that CRISPRa of *PKD1* can increase PC1 levels and suppress cellular features of PKD, including a decrease in intracellular cAMP levels, and an inhibition of cystogenic pathways and *in vitro* cyst formation. Given the remarkable renal plasticity and capacity for disease reversal previously observed upon *Pkd1* re-expression (36), our reagents offer a promising foundation for gene-augmentation strategies. While our short-term *in vitro* models demonstrate robust initial target engagement, evaluating the long-term durability of CRISPRa expression and whether repeat administration will be necessary, remains a critical step for future *in vivo* translational studies. Ultimately, optimizing these CRISPRa tools for *in vivo* delivery and human-relevant models will be crucial to translate this potential therapeutic approach into a transformative precision medicine for patients with ADPKD.

## Acknowledgments

We thank Dr. Ronak Lakhia (University of Texas Southwestern Medical Center) for providing the *Pkd1*^RC/-^ mouse collecting duct-derived epithelial cell line and Dr. Stephen Parnell (University of Kansas Medical Center-PKD Rodent Model and Drug-Testing Core) for providing primary mouse renal epithelial cells. We also acknowledge Drs. Hartmut Jaeschke, Anup Ramachandran, and Olamide Adelusi (University of Kansas Medical Center) for assistance with fluorescence microscopy. Finally, we thank Drs. Heather Wilkins and Ashley Tetlow at the University of Kansas Alzheimer’s Disease Research Center for their help with the Seahorse Assay and also the Flow Cytometry Core Laboratory at the University of Kansas Medical Center for technical assistance.

## Author Contributions

A.C. and A.S.L.Y. contributed to the study conception and design. A.C performed the experiments. M.M.V performed microcyst imaging. A.C and A.S.L.Y analyzed the data, prepared figures and wrote the manuscript. C.J.W developed and provided the anti-PC1 antibody. D.P.W provided primary human cells and along with C.J.W, edited and revised the manuscript. All authors read and approved of the final manuscript.

## References

1. Lanktree MB, Haghighi A, di Bari I, Song X, and Pei Y. Insights into autosomal dominant polycystic kidney disease from genetic studies. Clinical journal of the American Society of Nephrology 16: 790–799, 2021.

2. Gordon CE, Garimella PS, Perrone RD, and Miskulin DC. Autosomal dominant polycystic kidney disease: core curriculum 2025. American Journal of Kidney Diseases 2025.

3. Su Q, Hu F, Ge X, Lei J, Yu S, Wang T, Zhou Q, Mei C, and Shi Y. Structure of the human PKD1-PKD2 complex. Science 361: eaat9819, 2018.

4. Ong AC, and Harris PC. A polycystin-centric view of cyst formation and disease: the polycystins revisited. Kidney international 88: 699–710, 2015.

5. Qiu J, Germino GG, and Menezes LF. Mechanisms of cyst development in polycystic kidney disease. Advances in kidney disease and health 30: 209–219, 2023.

6. Lakhia R, Ramalingam H, Chang C-M, Cobo-Stark P, Biggers L, Flaten A, Alvarez J, Valencia T, Wallace DP, Lee EC, and Patel V. PKD1 and PKD2 mRNA cis-inhibition drives polycystic kidney disease progression. Nature communications 13: 4765, 2022.

7. Cheng AS, Li LX, Zhou JX, Harris PC, Calvet JP, and Li X. In vivo base editing rescues ADPKD in a humanized mouse model. Nature Communications 2025.

8. Vishy CE, Thomas C, Vincent T, Crawford DK, Goddeeris MM, and Freedman BS. Genetics of cystogenesis in base-edited human organoids reveal therapeutic strategies for polycystic kidney disease. Cell Stem Cell 31: 537–553. e535, 2024.

9. Rossetti S, Hopp K, Sikkink RA, Sundsbak JL, Lee YK, Kubly V, Eckloff BW, Ward CJ, Winearls CG, Torres VE, and Harris PC. Identification of gene mutations in autosomal dominant polycystic kidney disease through targeted resequencing. Journal of the American Society of Nephrology 23: 915–933, 2012.

10. Chavez A, Scheiman J, Vora S, Pruitt BW, Tuttle M, PR Iyer E, Lin S, Kiani S, Guzman CD, Wiegand DJ, Ter-Ovanesyan D, Braff JL, Davidsohn N, Housden BE, Perrimon N, Weiss R, Aach J, Collins JJ, and Church GM. Highly efficient Cas9-mediated transcriptional programming. Nature methods 12: 326–328, 2015.

11. Meylan P, Dreos R, Ambrosini G, Groux R, and Bucher P. EPD in 2020: enhanced data visualization and extension to ncRNA promoters. Nucleic acids research 48: D65–D69, 2020.

12. Concordet J-P, and Haeussler M. CRISPOR: intuitive guide selection for CRISPR/Cas9 genome editing experiments and screens. Nucleic acids research 46: W242–W245, 2018.

13. Lizio M, Harshbarger J, Shimoji H, Severin J, Kasukawa T, Sahin S, Abugessaisa I, Fukuda S, Hori F, Ishikawa-Kato S, Mungall CJ, Arner E, Baillie JK, Bertin N, Bono H, de Hoon M, Diehl AD, Dimont E, Freeman TC, Fujieda K, Hide W, Kaliyaperumal R, Katayama T, Lassmann T, Meehan TF, Nishikata K, Ono H, Rehli M, Sandelin A, Schultes EA, t Hoen PA, Tatum Z, Thompson M, Toyoda T, Wright DW, Daub CO, Itoh M, Carninci P, Hayashizaki Y, Forrest AR, Kawaji H, and consortium F. Gateways to the FANTOM5 promoter level mammalian expression atlas. Genome Biol 16: 22, 2015.

14. Bae S, Park J, and Kim J-S. Cas-OFFinder: a fast and versatile algorithm that searches for potential off-target sites of Cas9 RNA-guided endonucleases. Bioinformatics 30: 1473–1475, 2014.

15. Yates AD, Achuthan P, Akanni W, Allen J, Allen J, Alvarez-Jarreta J, Amode MR, Armean IM, Azov AG, Bennett R, Bhai J, Billis K, Boddu S, Marugan JC, Cummins C, Davidson C, Dodiya K, Fatima R, Gall A, Giron CG, Gil L, Grego T, Haggerty L, Haskell E, Hourlier T, Izuogu OG, Janacek SH, Juettemann T, Kay M, Lavidas I, Le T, Lemos D, Martinez JG, Maurel T, McDowall M, McMahon A, Mohanan S, Moore B, Nuhn M, Oheh DN, Parker A, Parton A, Patricio M, Sakthivel MP, Abdul Salam AI, Schmitt BM, Schuilenburg H, Sheppard D, Sycheva M, Szuba M, Taylor K, Thormann A, Threadgold G, Vullo A, Walts B, Winterbottom A, Zadissa A, Chakiachvili M, Flint B, Frankish A, Hunt SE, G II, Kostadima M, Langridge N, Loveland JE, Martin FJ, Morales J, Mudge JM, Muffato M, Perry E, Ruffier M, Trevanion SJ, Cunningham F, Howe KL, Zerbino DR, and Flicek P. Ensembl 2020. Nucleic Acids Res 48: D682–D688, 2020.

16. Smith TF, and Waterman MS. Identification of common molecular subsequences. Journal of molecular biology 147: 195–197, 1981.

17. Han Z, Madhavan BK, Kaymak S, Nawroth P, and Kumar V. A fast and reliable method to generate pure, single cell-derived clones of mammalian cells. Bio-protocol 12: e4490–e4490, 2022.

18. Yoshimura Y, Muto Y, Omachi K, Miner JH, and Humphreys BD. Elucidating the proximal tubule HNF4A gene regulatory network in human kidney organoids. Journal of the American Society of Nephrology 34: 1672–1686, 2023.

19. Schoger E, Carroll KJ, Iyer LM, McAnally JR, Tan W, Liu N, Noack C, Shomroni O, Salinas G, Gross J, Herzog N, Doroudgar S, Bassel-Duby R, Zimmermann WH, and Zelarayan LC. CRISPR-Mediated Activation of Endogenous Gene Expression in the Postnatal Heart. Circ Res 126: 6–24, 2020.

20. Chakraborty A, and Yu AS. Miniaturization of CRISPRa plasmids for efficient delivery into renal epithelial cells and Pkd1 transactivation. Molecular Biology Reports 53: 707, 2026.

21. Lea WA, Winklhofer T, Zelenchuk L, Sharma M, Rossol-Allison J, Fields TA, Reif G, Calvet JP, Bakeberg JL, Wallace DP, and Ward CJ. Polycystin-1 Interacting Protein-1 (CU062) Interacts with the Ectodomain of Polycystin-1 (PC1). Cells 12: 2023.

22. Schindelin J, Arganda-Carreras I, Frise E, Kaynig V, Longair M, Pietzsch T, Preibisch S, Rueden C, Saalfeld S, Schmid B, Tinevez JY, White DJ, Hartenstein V, Eliceiri K, Tomancak P, and Cardona A. Fiji: an open-source platform for biological-image analysis. Nat Methods 9: 676–682, 2012.

23. Hopp K, Ward CJ, Hommerding CJ, Nasr SH, Tuan H-F, Gainullin VG, Rossetti S, Torres VE, and Harris PC. Functional polycystin-1 dosage governs autosomal dominant polycystic kidney disease severity. The Journal of clinical investigation 122: 4257–4273, 2012.

24. Rowe I, Chiaravalli M, Mannella V, Ulisse V, Quilici G, Pema M, Song XW, Xu H, Mari S, Qian F, Pei Y, Musco G, and Boletta A. Defective glucose metabolism in polycystic kidney disease identifies a new therapeutic strategy. Nat Med 19: 488–493, 2013.

25. Podrini C, Rowe I, Pagliarini R, Costa ASH, Chiaravalli M, Di Meo I, Kim H, Distefano G, Tiranti V, Qian F, di Bernardo D, Frezza C, and Boletta A. Dissection of metabolic reprogramming in polycystic kidney disease reveals coordinated rewiring of bioenergetic pathways. Commun Biol 1: 194, 2018.

26. Scholz JK, Kraus A, Lüder D, Skoczynski K, Schiffer M, Grampp S, Schödel J, and Buchholz B. Loss of Polycystin-1 causes cAMP-dependent switch from tubule to cyst formation. Iscience 25: 2022.

27. Yamaguchi T, Wallace DP, Magenheimer BS, Hempson SJ, Grantham JJ, and Calvet JP. Calcium restriction allows cAMP activation of the B-Raf/ERK pathway, switching cells to a cAMP-dependent growth-stimulated phenotype. Journal of Biological Chemistry 279: 40419–40430, 2004.

28. Yamaguchi T, Nagao S, Wallace DP, Belibi FA, Cowley Jr BD, Pelling JC, and Grantham JJ. Cyclic AMP activates B-Raf and ERK in cyst epithelial cells from autosomal-dominant polycystic kidneys. Kidney international 63: 1983–1994, 2003.

29. Parrot C, Kurbegovic A, Yao G, Couillard M, Cote O, and Trudel M. C-MYC is a regulator ofthe PKD1 gene and PC1-induced pathogenesis. Hum Mol Genet 2018.

30. Gilbert LA, Horlbeck MA, Adamson B, Villalta JE, Chen Y, Whitehead EH, Guimaraes C, Panning B, Ploegh HL, Bassik MC, Qi LS, Kampmann M, and Weissman JS. Genome-Scale CRISPR-Mediated Control of Gene Repression and Activation. Cell 159: 647–661, 2014.

31. Konstantakos V, Nentidis A, Krithara A, and Paliouras G. CRISPR–Cas9 gRNA efficiency prediction: an overview of predictive tools and the role of deep learning. Nucleic Acids Research 50: 3616–3637, 2022.

32. Kemaladewi DU, Bassi PS, Erwood S, Al-Basha D, Gawlik KI, Lindsay K, Hyatt E, Kember R, Place KM, Marks RM, Durbeej M, Prescott SA, Ivakine EA, and Cohn RD. A mutation-independent approach for muscular dystrophy via upregulation of a modifier gene. Nature 572: 125–130, 2019.

33. Di Maria V, Moindrot M, Ryde M, Bono A, Quintino L, and Ledri M. Development and validation of CRISPR activator systems for overexpression of CB1 receptors in neurons. Frontiers in Molecular Neuroscience 13: 168, 2020.

34. Zhu H, Liu D, Sui M, Zhou M, Wang B, Qi Q, Wang T, Zhang G, Wan F, and Zhang B. CRISPRa-based activation of Fgf21 and Fndc5 ameliorates obesity by promoting adipocytes browning. Clin Transl Med 13: e1326, 2023.

35. Ibel A, Bhardwaj R, Yilmaz DE, Kong S, Wendlinger S, Cordero C, Papaioannou D, Papazian M, Schonauer R, Meng Q, Eckardt KU, Hassan F, Volpe I, Klambt V, Halbritter J, Fedeles S, Krappitz M, and Kaminski MM. In vivo base editing reduces liver cysts in autosomal dominant polycystic kidney disease. Mol Ther 33: 5373–5382, 2025.

36. Dong K, Zhang C, Tian X, Coman D, Hyder F, Ma M, and Somlo S. Renal plasticity revealed through reversal of polycystic kidney disease in mice. Nature genetics 53: 1649–1663, 2021.

37. Thivierge C, Kurbegovic A, Couillard M, Guillaume R, Coté O, and Trudel M. Overexpression of PKD1 causes polycystic kidney disease. Molecular and cellular biology 26: 1538–1548, 2006.

38. Patel V, Hajarnis S, Williams D, Hunter R, Huynh D, and Igarashi P. MicroRNAs regulate renal tubule maturation through modulation of Pkd1. Journal of the American Society of Nephrology 23: 1941–1948, 2012.

39. Erard N, Knott SR, and Hannon GJ. A CRISPR resource for individual, combinatorial, or multiplexed gene knockout. Molecular cell 67: 348–354. e344, 2017.

40. Gonatopoulos-Pournatzis T, Aregger M, Brown KR, Farhangmehr S, Braunschweig U, Ward HN, Ha KCH, Weiss A, Billmann M, Durbic T, Myers CL, Blencowe BJ, and Moffat J. Genetic interaction mapping and exon-resolution functional genomics with a hybrid Cas9-Cas12a platform. Nat Biotechnol 38: 638–648, 2020.

41. Tak YE, Horng JE, Perry NT, Schultz HT, Iyer S, Yao Q, Zou LS, Aryee MJ, Pinello L, and Joung JK. Augmenting and directing long-range CRISPR-mediated activation in human cells. Nature methods 18: 1075–1081, 2021.

42. Casas-Mollano JA, Zinselmeier MH, Erickson SE, and Smanski MJ. CRISPR-Cas activators for engineering gene expression in higher eukaryotes. The CRISPR journal 3: 350–364, 2020.

43. Chen M, and Qi LS. Repurposing CRISPR system for transcriptional activation. RNA Activation 147–157, 2017.

