## Supplementary Information for "CRISPR activation of endogenous *PKD1* increases polycystin-1 levels and suppresses cellular features of ADPKD"

### Supplemental Tables

Suppl. Table S1. Summary of the cell lines used for CRISPRa experiments

Suppl. Table S2. Human *PKD1* guides

Suppl. Table S3. Mouse *Pkd1* guides

Suppl. Table S4. Primers

### Supplemental Figures

Suppl. Fig. S1. Promoter-reporter plasmid maps

Suppl. Fig. S2. Validation of HEK293T cells constitutively expressing dCas9-VPR

Suppl. Fig. S3 Validation of M1 cells constitutively expressing dCas9-VPR

Suppl. Fig. S4 Off-target validation in *Pkd1*$^{RC/-}$ cell model

Suppl. Fig. S5. Off-target validation in primary mouse PKD cells

Suppl. Fig. S6 .Off-target validation and *PKD1* promoter GC content

Suppl. Fig. S7. Pairwise and multiple sequence alignment

**Suppl. Table S1.** Summary of the cell lines used for CRISPRa experiments

| Cell line | Immortalized | Details | Additional transgenes[1] | Assay/purpose[2] |
|---|---|---|---|---|
| HEK-dCas9-VPR | Yes | Human cell line stably expressing dCas9-VPR | Human or mouse *PKD1*-EGFP reporter | Fluorescence screening for optimal human and mouse sgRNAs |
| | | | - | Endogenous human *PKD1* mRNA expression |
| M1-dCas9-VPR | Yes | Mouse collecting duct cell line stably expressing dCas9-VPR | - | Endogenous mouse *Pkd1* mRNA expression |
| *Pkd1*$^{RC/-}$ cell line | Yes | Mouse collecting duct cell line with single hypomorphic *Pkd*1 RC allele | mdCas9-VPR | Endogenous mouse *Pkd1* (RC) mRNA and polycystin-1 protein expression<br>Cell proliferation, 3D cyst growth, mitochondrial function, signaling assays |
| *Pkd1*$^{RC/Cond}$*; Pkhd1*$^{Cre-}$ primary cells i.e *RC/-* cells | No | Primary cells isolated from kidneys of mice with single hypomorphic *Pkd1* (RC) allele | - | Endogenous mouse *Pkd1* (RC) mRNA expression<br>Cell proliferation assay |
| NHK primary cells | No | Primary cells isolated from normal human kidneys | - | Endogenous human *PKD1* mRNA expression |
| Human ADPKD primary cells | No | Primary cells isolated from cyst epithelium of human ADPKD nephrectomy specimens | - | Endogenous human *PKD1* mRNA expression |

[1]Additional genes transiently transfected into cells. All cells were also transiently (co-)transfected with sgRNA targeted to the *PKD1* gene for the appropriate species (mouse or human) or a negative control ("non-targeting") sgRNA

[2]HEK-dCas9-VPR was used as an all-purpose dCas9-VPR-expressing cell line for screening of both human and mouse sgRNAs using an exogenous *PKD1*-EGFP reporter. HEK dCas9-VPR was also used to evaluate the ability of human PKD1-targeted sgRNAs to activate its endogenous human *PKD1* gene expression, while the mouse M1-dCas9-VPR cell line was used to evaluate mouse sgRNAs for activation of endogenous mouse *Pkd1* gene expression. To evaluate the functional effect of *Pkd1* transcriptional activation, assays were performed in an immortalized mouse cell line with a hypomorphic RC allele (*Pkd1*$^{RC/-}$). Some of these assays were confirmed in primary cells isolated from kidneys of mice with the same genotype. Finally, human sgRNAs were tested for transcriptional activation in human primary cells from both normal kidneys (NHK) and ADPKD cyst epithelial cells. Note that many assays were not feasible in primary cells due to their technical challenges with efficient transfection, cell number and availability.

**Suppl. Table S2.** Human *PKD1* guides

| sgRNA ID | Guide sequence (5'- 3') | PAM | Strand | Chromosomal location | Position relative to TSS (bp) |
|---|---|---|---|---|---|
| sgRNA1 | GAGGAGGAGGAGCCGCGGCG | GGG | - | Chr16:2135905-2135924 | -28 |
| sgRNA2* | GCGCGCGGCGCGGGGCGGAC | GGG | + | Chr16:2135929-2135948 | -33 |
| sgRNA3* | GTCCTCCTCCCCGCGCGGCG | CGG | + | Chr16:2135919-2135938 | -23 |
| sgRNA4* | GGGGGGGGCGGGGCGGGTGC | AGG | + | Chr16:2135955-2135974 | -59 |
| sgRNA9* | GGCCCGTCGCACTGCAGAGT | CGG | + | Chr16:2136142-2136161 | -246 |
| sgRNA10* | GACTTTAGCCTGCAGCGGGG | CGG | - | Chr16:2136194-2136213 | -319 |
| sgRNA12 | GGCTTCACCCTCCGCTCCAC | AGG | + | Chr16:2136034-2136053 | -138 |
| sgRNA15 | GGATGCCAGTCCCTCATCGC | TGG | - | Chr16:2136011-2136030 | -134 |
| sgRNA16* | GGCAGCGCCAGCGTCCGAGC | GGG | - | Chr16:2135875-2135894 | +1 |
| sgRNA17 | GCCTGGCCCCGAGCCCCGAG | CGG | - | Chr16:2135833-2135852 | +44 |
| sgRNA18* | GATGCGCCCCGGGAACGCGT | GGG | - | Chr16:2136094-2136113 | -217 |

*These sgRNAs were modified by replacing the first 5' nucleotide with a guanine (G). The U6 promoter, used to express the sgRNA, is an RNA polymerase III promoter that strongly prefers a G to initiate transcription. Adding a G, particularly when the spacer sequence does not begin with one, improves the transcriptional efficiency of the sgRNA.

**Suppl. Table S3.** Mouse *Pkd1* guides

| sgRNA ID | Guide sequence (5'- 3') | PAM | Strand | Chromosomal location | Position relative to TSS (bp)[#] |
|---|---|---|---|---|---|
| sgRNA1* | GATAAGCTCTCCAGCCGTTG | TGG | + | Chr17: 24549646 - 24549665 | -408 |
| sgRNA2 | GCCCCGCCTTCTGCACCGCT | GGG | - | Chr17: 24549785 - 24549804 | -249 |
| sgRNA3* | GACCAAGGCTCACGCTCACT | GGG | + | Chr17: 24549838 - 24549857 | -216 |
| sgRNA4* | GCGGGAGGGGAGTCCTCCGG | AGG | - | Chr17: 24549807 - 2135974 | -228 |
| sgRNA5 | GGCGTTTCGAGCCCCGAGTC | AGG | - | Chr17: 24549711 - 24549730 | -324 |
| sgRNA6* | GGTCCCCTACCCACAACGGC | TGG | - | Chr17: 24549656- 24549678 | -376 |
| sgRNA7* | GTCACTGGGCGGAGCCTCTG | AGG | + | Chr17:2 24549852 - 24549871 | -202 |
| sgRNA8* | GGAAGGAAGGGCGCCCTCAG | AGG | - | Chr17: 24549869 - 24549888 | -166 |
| sgRNA9* | GTTGGTGGGCGGGGTCTCAC | GGG | - | Chr17: 24549825 - 24549844 | -210 |
| sgRNA10 | GTGGGCGGGGTCTCACGGGA | GGG | - | Chr17: 24549821 - 2135852 | -214 |
| sgRNA11* | GACAGCCAACTTGGAAGCGC | AGG | + | Chr17: 24549962 - 24549981 | -93 |
| sgRNA12* | GCGCCCCTGCGCTTCCAAGT | TGG | - | Chr17: 24549970 - 24549989 | -65 |
| sgRNA13* | GCCTCCCCCACCTCCGGCGT | GGG | - | Chr17: 24550011 - 24550030 | -24 |
| sgRNA14* | GGCAGCGCGAATGCGCGAGC | AGG | + | Chr17: 24550056 - 24550075 | +2 |
| sgRNA15* | GATGCGCGAGCAGGCGGCCA | AGG | + | Chr17: 24550065 - 24550084 | +11 |
| NT-sgRNA* | GAGGAGTCGCCGATACGCGT | - | - | - | - |

*These sgRNAs were modified by replacing the first 5' nucleotide with a guanine (G).

[#]The mm10 FANTOM5 CAGE atlas identifies four putative TSS for mouse *Pkd1*, whereas the Eukaryotic Promoter Database identifies only one. To maintain consistency, sgRNA distances were measured from the TSS shared by both datasets.

**Suppl. Table S4.** Primers

| Target | Forward sequence (5' – 3') | Reverse sequence (5' – 3') |
|---|---|---|
| *dCas9* | AGACACGCCAGATCACCAAG | TCGATAAGTGGTCGCTTCCG |
| *HNF4A* | CCTACCTCAAAGCCATCAT | ATGTAGTCCTCCAAGCTCAC |
| *PKD1* | CGGCTGCAGGAAGCACTCTA | AACGTCGTAATCGCTGGTGC |
| *RAB26* | GACGTCGCCTTCAAGGTCAT | ATCCTTGAATCGCACCAGCA |
| *GAPDH* | TCGGAGTCAACGGATTTGGT | TGAAGGGGTCATTGATGGCA |
| *Klf15* | CAACTCATCTGAGCGGGAA | CAAGAGCAGCCACCTCAAG |
| *Pkd1* | GATCAGACACCGCTCAACTTCC | ACACCAGCTTCTAGGCGTTCCA |
| *Sart3* | ATTACAACCTGGAACGGGCA | CACAGACGTGCTCAGGGTAG |
| *Cds1* | TCATATCCTTCGCCCTCTACC | GCGTCCATGCGAACATATAGA |
| *Rab26* | GATGTTGCTTTCAAGGTCATGCT | ATGCCCACAGTGGAGATGAAG |
| *Gapdh* | GGGTCCCAGCTTAGGTTCATCAG | ATCCGTTCACACCGACCTTCAC |

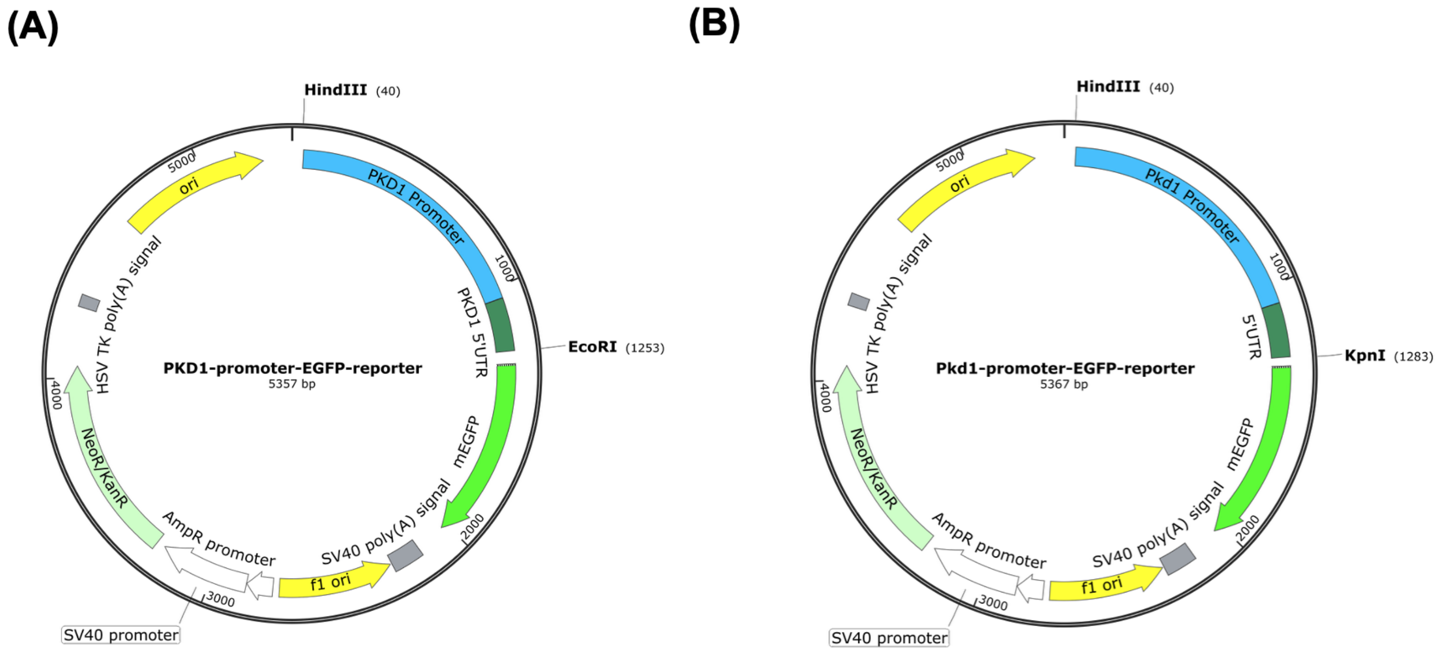


**Supplemental Fig. S1**. Promoter-reporter plasmid maps. **(A)** Plasmid map of the EGFP reporter fused to the 5′ UTR and 1,000 bp promoter regions of human *PKD1* (ENSG00000008710) upstream of the TSS. **(B)** Plasmid map of the EGFP reporter fused to the 5′ UTR and 1,000 bp promoter regions of mouse *Pkd1* (ENSMUSG00000032855) upstream of the TSS. The nucleotide sequences were retrieved from the Ensembl database 100. EGFP (light green) = Enhanced green fluorescent protein, UTR (dark green) = Untranslated region, TSS = Transcription start site

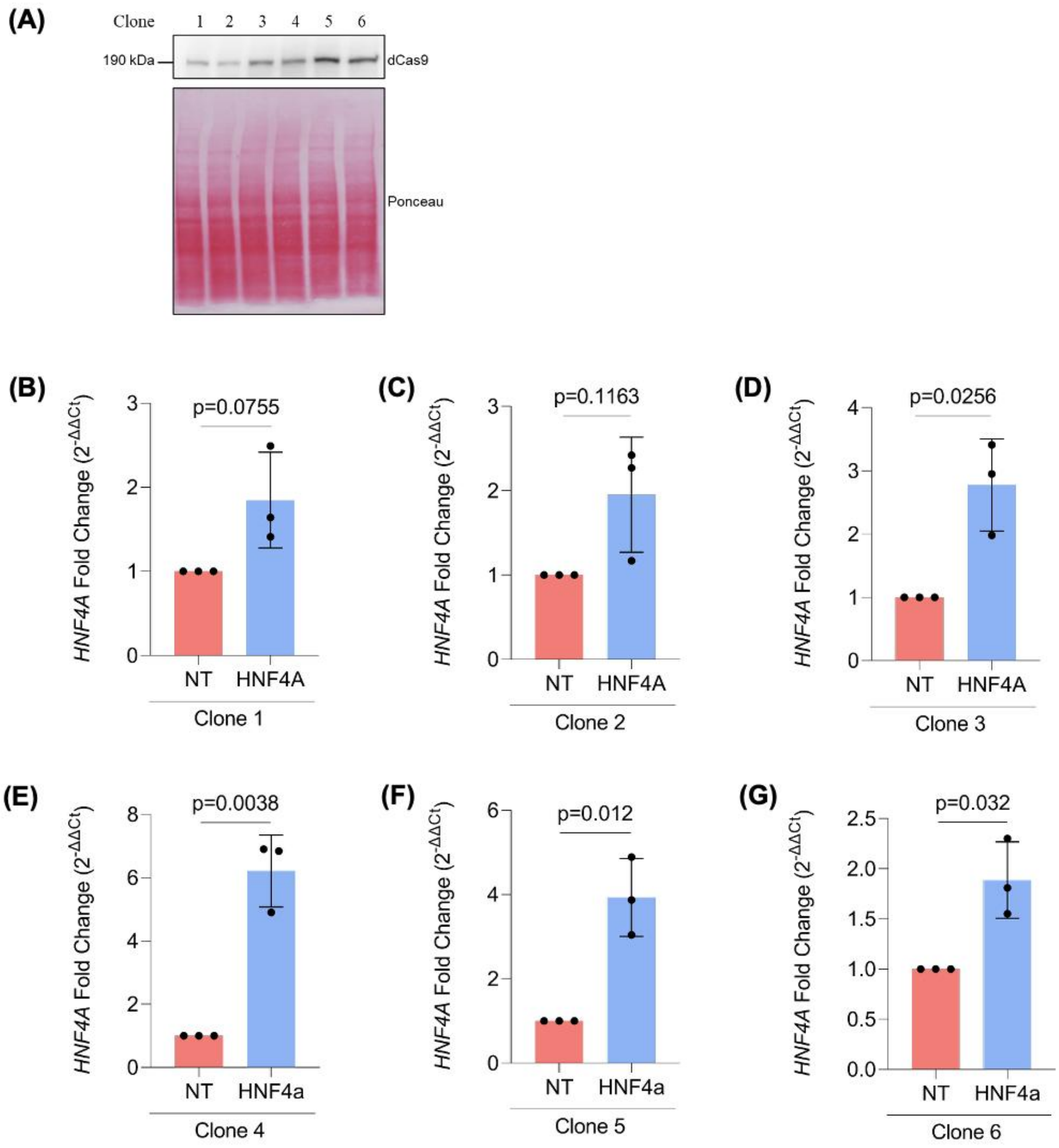


**Supplemental Fig. S2.** Validation of HEK293T cells constitutively expressing dCas9-VPR. **(A)** Immunoblot of dCas9 identified by an anti-Cas9 antibody in HEK-dCas9-VPR clones 1-6. **(B-G)** *HNF4A* mRNA expression in HEK-dCas9-VPR clones 1-6 following transfection with either NT-sgRNA or *HNF4A*-targeting-sgRNA. Data are expressed as fold change relative to the NT-sgRNA (N=3 biological replicates; mean ± SD). Statistical significance was determined by a one-sample *t*-test performed on -ΔΔCq values compared to a theoretical mean of 0.

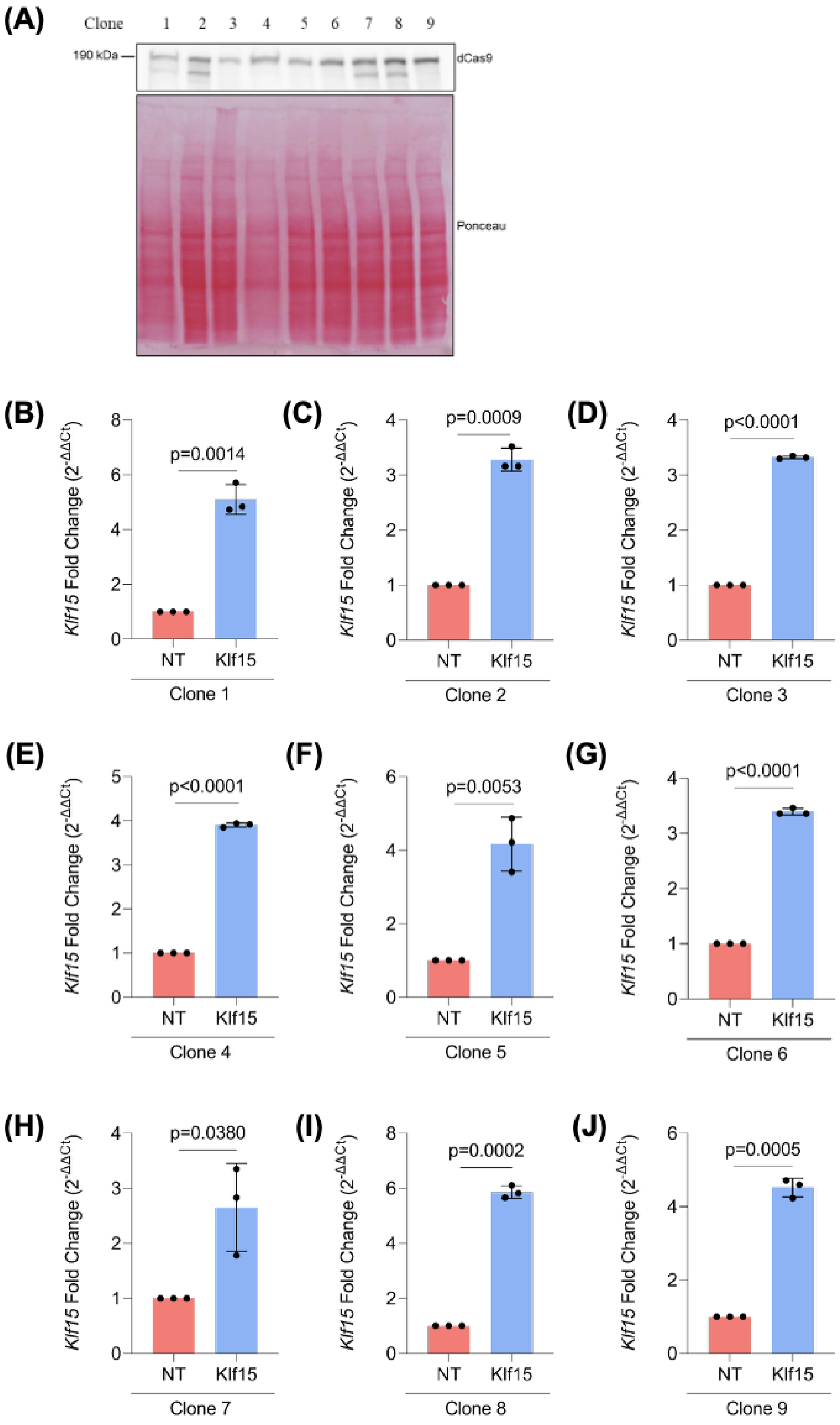


**Supplemental Fig. S3.** Validation of M1 cells constitutively expressing dCas9-VPR. **(A)** Immunoblot of dCas9 identified by an anti-Cas9 antibody in M1-dCas9-VPR clones 1-9. **(B-J)** Relative *Klf15* mRNA expression in M1-dCas9-VPR clones 1-9 following transduction with either NT-sgRNA or *Klf15*-targeting-sgRNA. Data are expressed as fold change relative to the NT-sgRNA (N=3 biological replicates; mean ± SD). Statistical significance was determined by a one-sample *t*-test performed on -ΔΔCq values compared to a theoretical mean of 0.

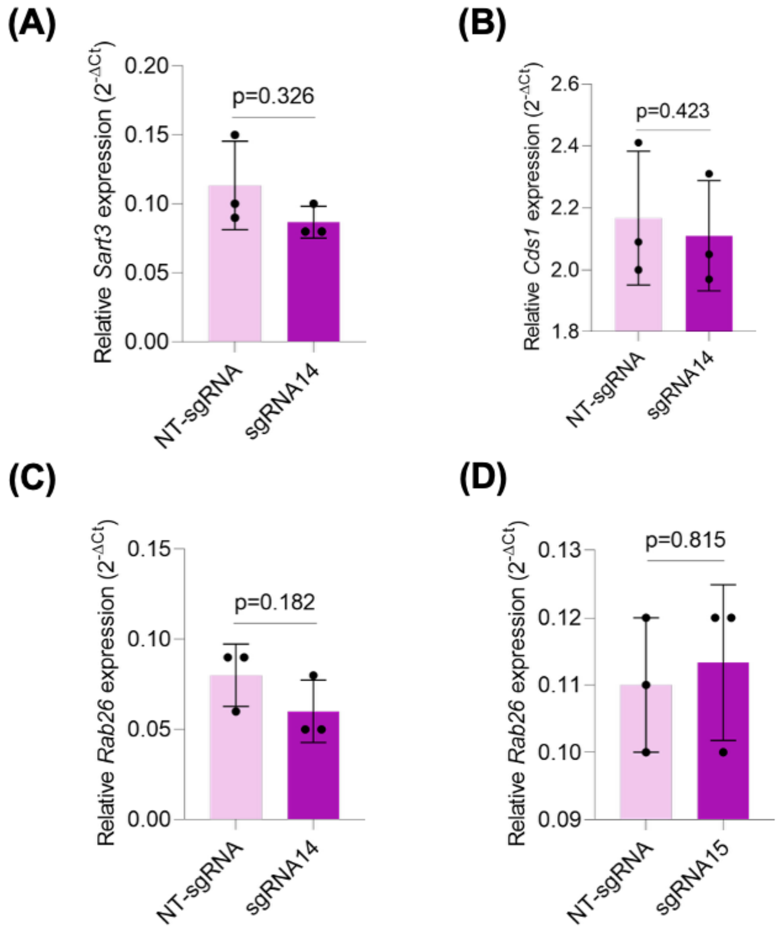


**Supplemental Fig. S4.** Off-target validation in *Pkd1*$^{RC/-}$ cell model. **(A-D)** Relative transcript levels of **(A)** *Sart3* **(B)** *Cds1* and **(C-D)** *Rab26* in primary *Pkd1*$^{RC/-}$ cells following nucleofection with mdCas9-VPR and indicated sgRNAs. For all panels**,** data are presented as mean $2^{-\Delta Cq}$ ± SD (N=3 biological replicates). Statistical significance was determined by a two-tailed paired *t*-test on -ΔCq values.

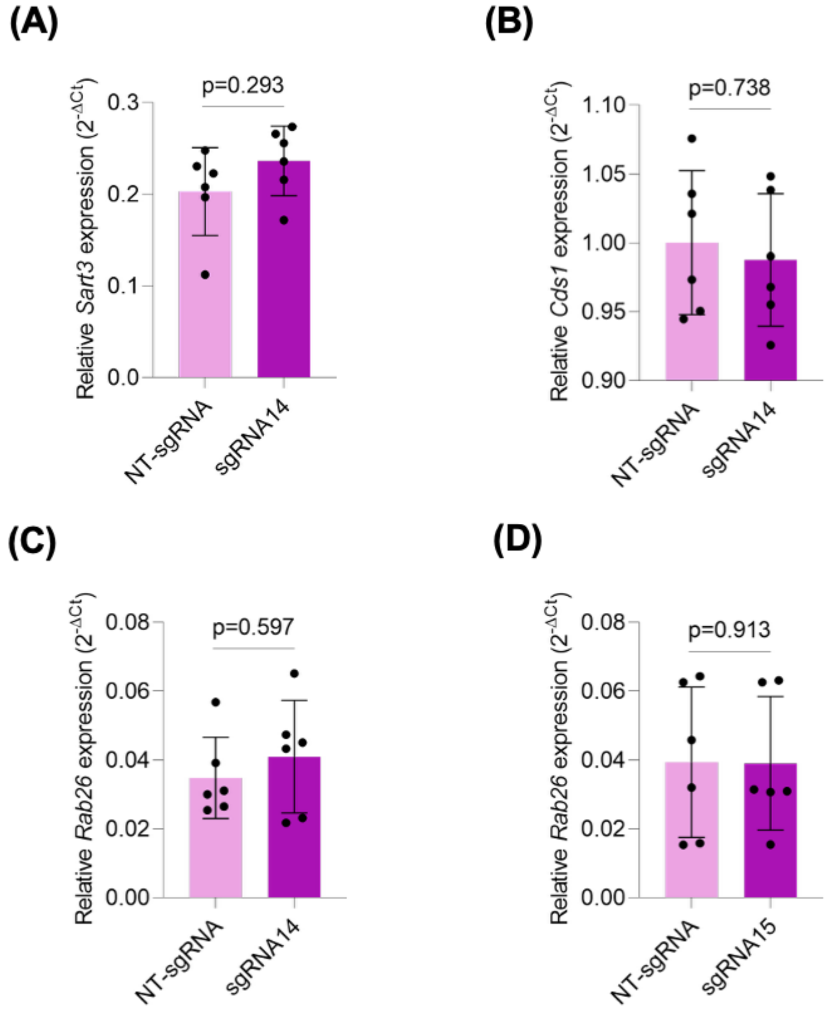


**Supplemental Fig. S5.** Off-target validation in primary mouse PKD cells. **(A-D)** Relative transcript levels of **(A)** *Sart3* **(B)** *Cds1* and **(C-D)** *Rab26* in primary *RC/-* cells ($Pkd1^{RC/Cond}$; $Pkhd1^{Cre+}$) cells following nucleofection with mdCas9-VPR and indicated sgRNAs. For all panels, data are presented as mean $2^{-\Delta Cq}$ ± SD (N=3 biological replicates). Statistical significance was determined by a two-tailed paired *t*-test on -ΔCq values.

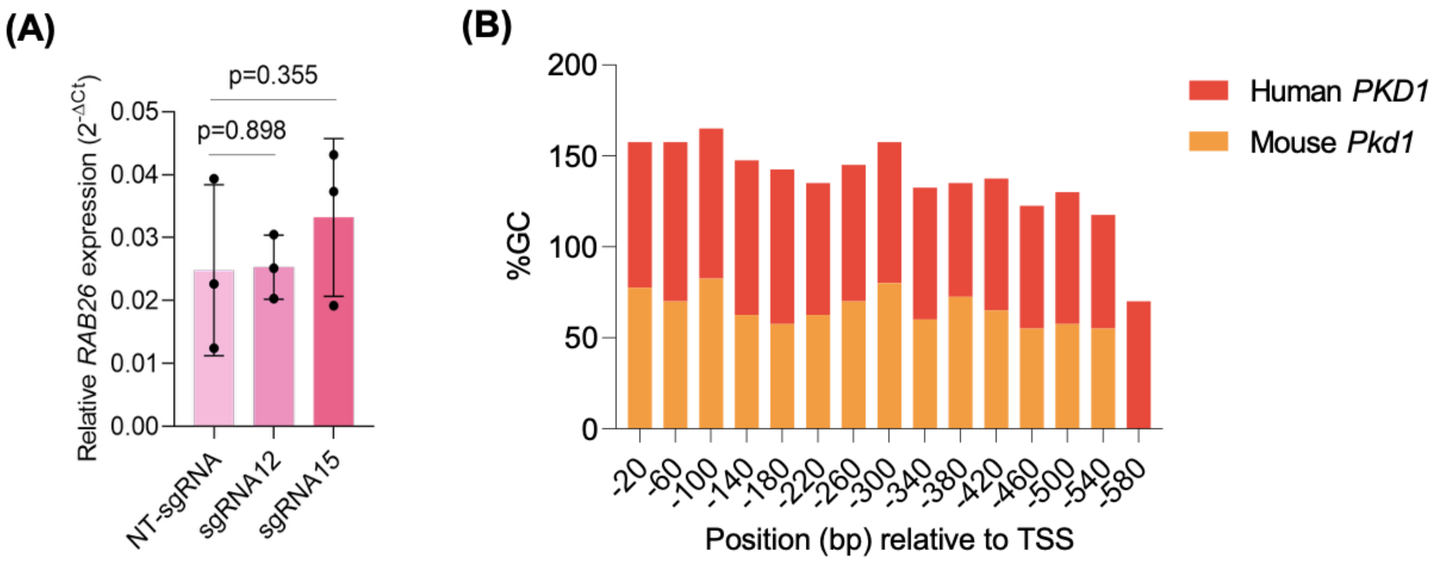


**Supplemental Fig. S6.** Off-target validation and *PKD1* promoter GC content. **(A)** Relative *RAB26* transcript levels in primary NHK cells following nucleofection with mdCas9-VPR and indicated sgRNAs to assess potential bystander activation. Data are presented as mean $2^{-\Delta Cq}$ ± SD (N = 3). Statistical significance was determined by repeated measures one-way ANOVA on -ΔCq values followed by Dunnett's multiple comparisons test. **(B)** GC content (%) was calculated using a 40 bp sliding window across the genomic regions targeted by the mouse and human-sgRNAs, spanning approximately 600 bp upstream of the transcription start site (TSS). Data are presented as a stacked bar graph to compare regional GC distribution between human *PKD1* (red) and mouse *Pkd1* (orange).

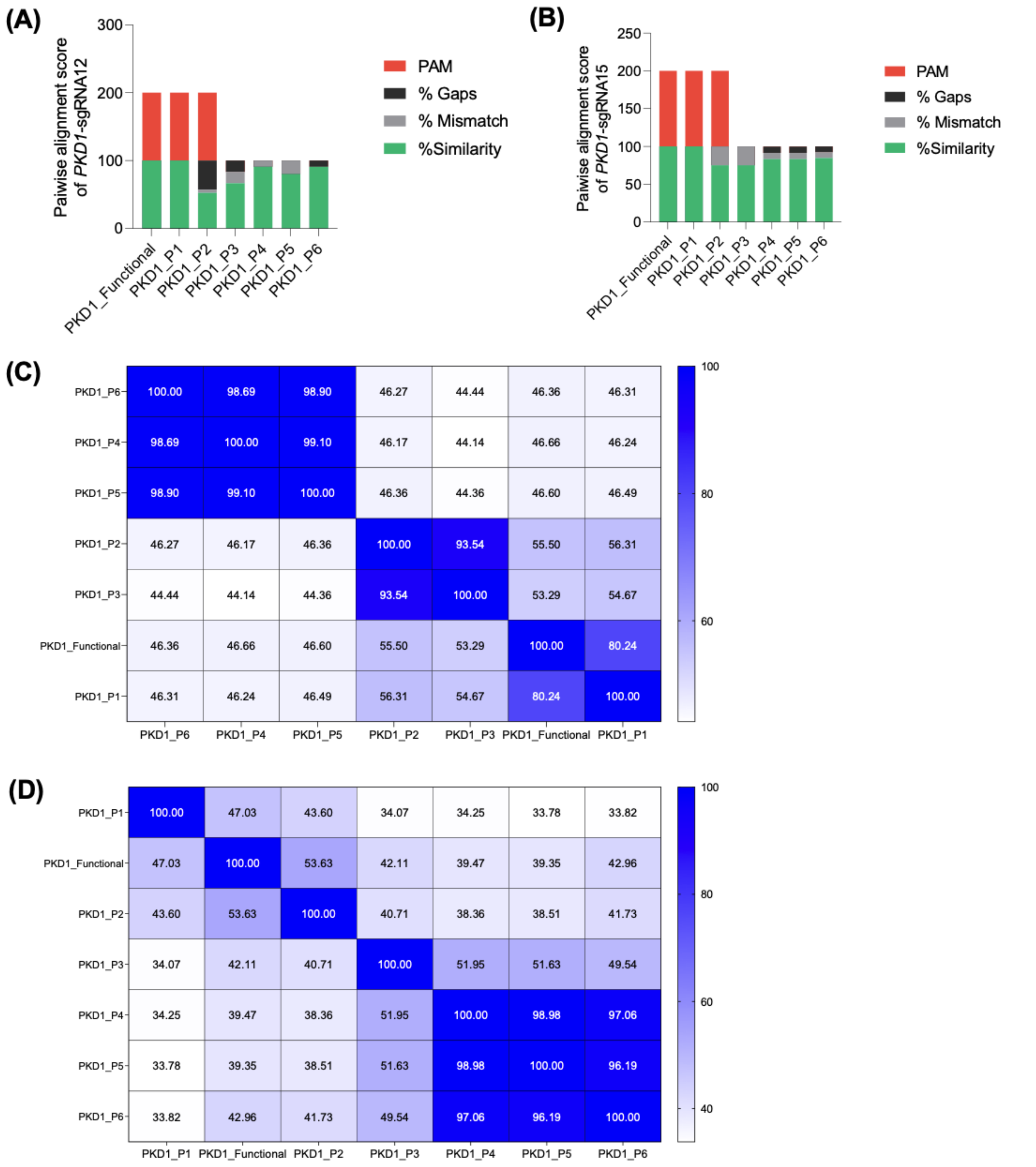


**Supplemental Fig. S7.** Pairwise and multiple sequence alignment. **(A-B)** Pairwise alignment of the 12 bp seed sequence of human **(A)** *PKD1*-sgRNA12 and **(B)** *PKD1*-sgRNA15 with the promoter regions of the functional *PKD1* gene and its six paralogs (*PKD1P1-P6)* using EMBOSS Water. The Smith-Waterman algorithm was used to calculate percent similarity, mismatch frequency, and gap occurrence. The presence or absence of a 'NGG' PAM was scored as 100 or 0, respectively. **(C-D)** Heatmap showing the percent sequence identity of the **(C)** 2 kb and **(D)** 200 bp sequences upstream of the transcription start site between the functional *PKD1* and its pseudogenes as determined by Clustal Omega Multiple Sequence Alignment. Color scale indicates percent identity from low (cool blue) to high (warm blue).
